# Gating of hair cell Ca^2+^ channels governs the activity of cochlear neurons

**DOI:** 10.1101/2025.01.23.634330

**Authors:** Nare Karagulyan, Anupriya Thirumalai, Susann Michanski, Yumeng Qi, Qinghua Fang, Haoyu Wang, Nadine J. Ortner, Jörg Striessnig, Nicola Strenzke, Carolin Wichmann, Yunfeng Hua, Tobias Moser

## Abstract

Our sense of hearing processes sound intensities spanning six orders of magnitude. In the ear, postsynaptic spiral ganglion neurons (SGNs) tile this intensity range with their firing rate codes. Presynaptic inner hair cells (IHCs) vary Ca^2+^-influx among their active zones (AZs) diversifying glutamate release and likely contributing to SGN firing diversity. Here we show that low-voltage activation of IHC-Ca^2+^-influx of mice, modeling the human Ca_V_1.3^A749G^ mutation, increases spontaneous SGN-firing and lowers sound threshold. Altered synaptic morphology in Ca_V_1.3^A749G/A749G^ mice already at ambient sound levels of standard mouse husbandry indicates a risk for noise-induced alterations in Ca_V_1.3^A749G^ patients. We conclude that heterogeneous voltage-dependence of Ca_V_1.3 activation among IHC-AZs contributes to the diversity of SGN firing for sound intensity coding and synaptic vulnerability.

## Main Text

IHCs sample sound information at each tonotopic position of the cochlea and convey it to SGNs via afferent ribbon synapses that vary in structure and function. Such synaptic heterogeneity might contribute to diverse SGN responses that enable a tiling of the audible range of sound pressures by the rate codes of the individual SGNs. This collective intensity coding by functionally diverse SGNs likely supports high fidelity processing across the 120 decibel range (six orders of magnitude) of audible sound pressures (*1*). A current challenge is to causally relate synaptic heterogeneity (*2–9*) to the diverse molecular profiles (*10–13*) and physiology (*14–19*) of SGNs. For the interpretation of findings regarding SGN function, the field has embarked on the concept of a spatial segregation of SGN synapses on the basal pole of the IHC. SGNs with low spontaneous firing rate (SR) and high sound threshold (“low SR”) preferentially synapse on the “modiolar” IHC side (facing the modiolus, i.e., cochlear center), while high SR, low threshold (“high SR”) SGNs tend to innervate the opposite “pillar” side of the IHC (facing the pillar cells) (*6*, *20*). For example, the molecularly defined type I_b_ and I_c_ SGNs tend to innervate the modiolar side and, hence, have been considered to represent low SR SGNs, e.g., due to different excitability (*10*, *11*, *19*, *21*, *22*).

Similarly, the finding that pillar synapses activate at lower voltages than modiolar ones has sparked the hypothesis that differences in the voltage-range of synapse operation contribute to the variance in spontaneous and sound evoked firing among SGNs at a given tonotopic position (*2*, *4*). This presynaptic hypothesis is centered on differences in the voltage-dependence of Ca_V_1.3 Ca^2+^ channel activation among the AZ of a given IHC (*2*, *4*). Bridging gaps between the different approaches, elucidating and relating the likely intertwining candidate mechanisms will be key to creating a unified concept on sound intensity coding in the cochlea (*1*). In a previous study, mouse mutants for the putative Ca_V_1.3 regulator, *Gipc3*, showed a shift in the activation of synaptic Ca^2+^ influx to lower voltages with concomitant increase in SGN spontaneous firing (*2*). However, pillar synapses in IHCs of *Gipc3* mutant mice combined negative activation with increased maximal Ca^2+^ influx and IHC mechanotransduction was impaired, which complicates the interpretation of how synaptic transfer and SGN activity are related. This calls for manipulation of Ca_V_1.3 channel function not afflicted with potentially confounding changes in IHC physiology. Here, we capitalized on Ca_V_1.3^A749G^ (or *Cacna1d* ^A749G^) mouse mutants (*23*) modeling the human *CACNA1D* p.A749G mutation (*24*). While loss of Ca_V_1.3 function causes syndromic deafness (*25*), the *CACNA1D* p.A749G mutation belongs to a group of mutations that cause aberrant gating and often are associated with neurodevelopmental disorders with and without endocrine symptoms (*26*). The mutation shifts the Ca_V_1.3 activation to lower voltages which could provide a unique opportunity to test the presynaptic hypothesis of functional SGN diversity. However, as it also shifts inactivation to lower voltages (*24*), the Ca_V_1.3^A749G^ mutation might also deplete activatable Ca^2+^ channels of IHCs that are thought to rest at a potential -58 mV in the absence of sound stimulation *in vivo* (*27*). In fact, enhanced Ca_V_1.3 inactivation due to lack of Ca^2+^ binding protein(s) 1 and 2 impair hearing by reducing the number of activatable Ca_V_1.3 channels mediating synaptic sound encoding (*28*, *29*), with *CABP2* being another deafness gene (*30*). Hence, it was conceivable that hetero- and homozygous Ca_V_1.3^A749G^ mice (for simplicity we nicknamed the mutant allele Ca_V_1.3^AG^) and the (heterozygous) Ca_V_1.3^AG^ patients could either show supernatural acoustic sensitivity or, in case of excessive Ca_V_1.3 inactivation, be hearing impaired. Finally, Ca_V_1.3 gain-of-function might lead to noise-induced synaptic damage or synaptopathy (*31–33*) even at otherwise non-damaging noise levels. This, again, would inform clinical care, and potentially recommend noise protection, for patients affected by Ca_V_1.3^AG^ or other *CACNA1D* mutations that suffer from severe neurodevelopmental disorder (*24*) and, therefore, have not yet been analyzed for a potential auditory phenotype to our knowledge.

Here we employed a multidisciplinary approach to characterize the auditory system of Ca_V_1.3^AG/WT^ mice, modeling the human neurodevelopmental disorder (*24*), and Ca_V_1.3^AG/AG^ mice to leverage the maximal impact of the mutation on sound encoding. First, we analyzed voltage-gated Ca^2+^ currents of IHCs at the whole-cell and synaptic levels in apical cochlear coils, acutely dissected from hearing mice (postnatal days (p) 21-28). Maximal Ca^2+^ current amplitudes of Ca_V_1.3^AG/WT^ and Ca_V_1.3^AG/AG^ IHCs (fig. S1A and B) were not significantly different from control (Ca_V_1.3^WT/WT^), similar to what was observed in adrenal chromaffin cells of Ca_V_1.3^AG/WT^ mice (*23*), but contrasting the increased amplitudes found upon heterologous expression of the mutant Ca_V_1.3 channel (*24*). However, in keeping with heterologous expression (*24*), we found a gene-dose dependent shift of Ca^2+^ channel activation to lower voltages and increased voltage-sensitivity of activation in Ca_V_1.3^AG/AG^ IHCs (fig. S1C). Alike Ca^2+^ influx, exocytosis of Ca_V_1.3^AG/AG^ assessed as change in membrane capacitance (ΔC_m_) was activated at lower voltage (fig. S1D). When adjusting depolarizations to the potential of maximal Ca^2+^ influx in Ca_V_1.3^AG/AG^ IHCs (-37 mV vs. -17 mV in Ca_V_1.3^WT/WT^), we found IHC exocytosis largely unaltered (fig. S1E). The kinetics of activation and inactivation of Ca^2+^ channels in IHCs of Ca_V_1.3^AG/WT^ and Ca_V_1.3^AG/AG^ mice were unaltered, but, as found with heterologous expression (*24*) deactivation was slowed significantly in IHCs of both genotypes suggesting a higher prevalence of longer open times (mode 2 (*34*), fig. S2). Indeed, non-stationary fluctuation analysis of IHC Ca^2+^ influx (*35*, *36*) revealed an increase in the open probability which was balanced by a decrease of the activatable number of Ca^2+^ channels in Ca_V_1.3^AG/AG^ IHCs (fig. S3). Next, we tested whether this decrease reflects inactivation or a potential homeostatic reduction of the Ca_V_1.3 channels. Performing immunohistochemistry, we revealed a reduced number of Ca_V_1.3 channels at AZs of Ca_V_1.3^AG/AG^ IHCs evident as lower Ca_V_1.3 immunofluorescence intensity in confocal microscopy (fig. S4A and C) and reduced size of Ca_V_1.3 channel clusters in STED nanoscopy (fig. S4D and E). Likewise, the Ribeye/Ctbp2 immunofluorescence intensity was reduced indicating a concomitant decrease of ribbon size in apical IHCs (fig S4B).

We then employed spinning disc confocal Ca^2+^ imaging to analyze Ca^2+^ channel function at single IHC AZs (*2*, *4*): We identified single AZs by labeling their ribbons by a Ribeye/Ctbp2-binding fluorescently labeled peptide (*37*, *38*) and then recorded the change in fluorescence of the green Ca^2+^ indicator Fluo4-FF (k_D_: 10 µM, Fig. 2A). In keeping with the reduction of Ca_V_1.3 channel number of IHC AZs (fig. S4), we found a reduction of the maximal amplitude of the Ca^2+^ signals (ΔF/F_0 max_, approximating the maximal AZ Ca^2+^ influx (*37*)) of Ca_V_1.3^AG/AG^ AZs (Fig. 1B and Bi). AZs of Ca_V_1.3^AG/AG^ IHCs showed a -15 mV shift of the voltage of half-maximal activation (Fig. 1C and Cii) and increased voltage sensitivity of Ca^2+^ influx activation (Fig. 1Ci) which confirmed the notion of the whole IHC recording for the average single AZ. Next, we compared the properties of synapses of the modiolar and pillar IHC sides and found the modiolar-pillar gradient of maximal AZ Ca^2+^ influx (greater ΔF/F_0 max_ for modiolar synapses) typical for Ca_V_1.3^WT/WT^ IHCs (*2*) to be collapsed in Ca_V_1.3^AG/AG^ IHCs (Fig. 1D). This finding was contrasted by differences in immunofluorescence intensity for Ca_V_1.3 of modiolar and pillar AZs (fig. S5), which together with Ribeye/Ctbp2 immunofluorescence indicated that Ca_V_1.3^AG/AG^ IHCs maintain larger ribbons with greater complement of Ca_V_1.3 for modiolar AZ. Comparing the voltage operating range of AZs showed that Ca_V_1.3^AG/AG^ IHCs retained a significant pillar-modiolar gradient (lower V_half_ at pillar synapses (*2*), Fig. 1E). This was also evident from correlations of V_half_ and synapse position along the pillar-modiolar axis (fig. S6).

**Fig. 1.**
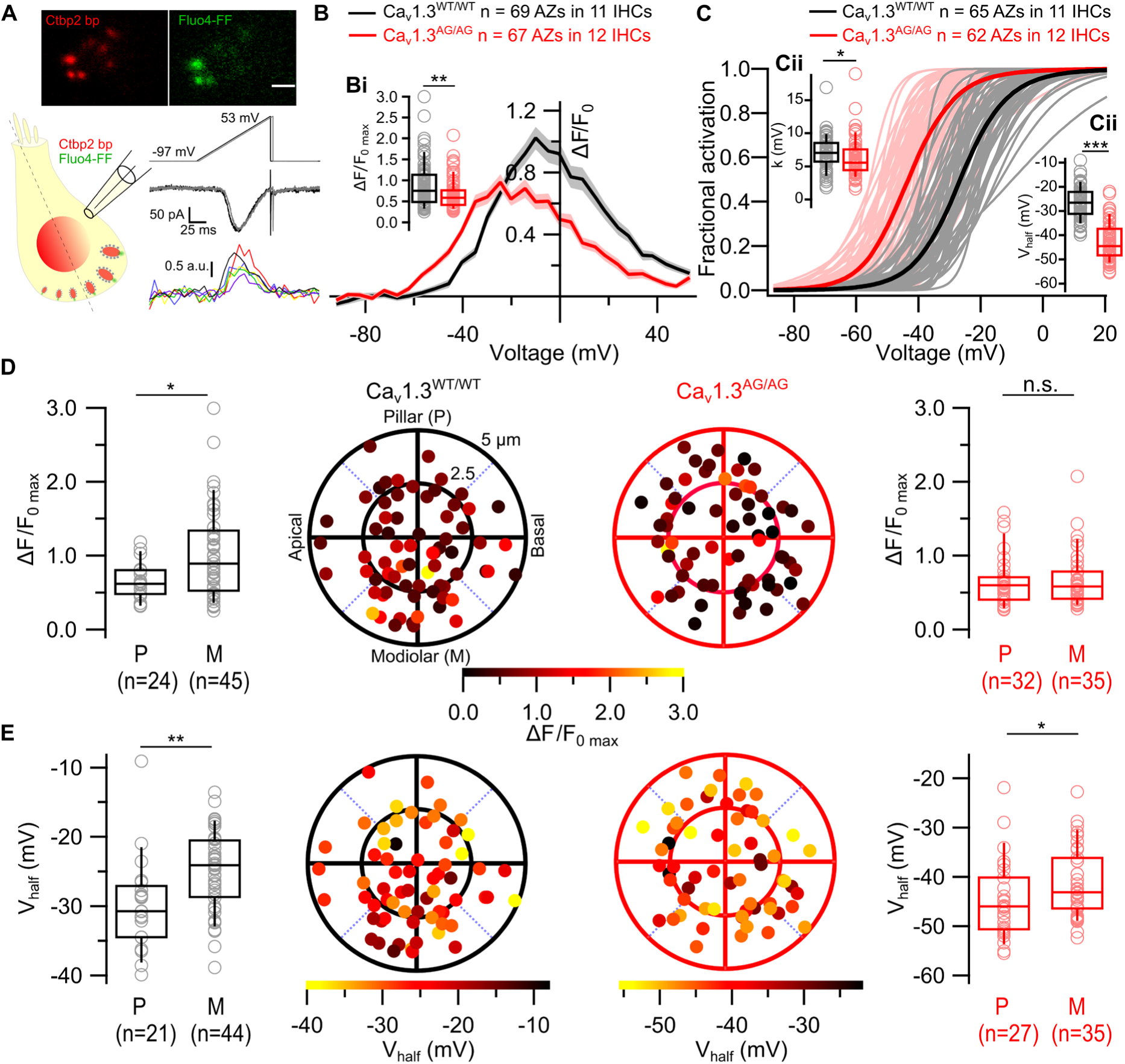
Reduced amplitude, hyperpolarized activation and altered voltage sensitivity of synaptic Ca^2+^ influx in Ca_V_1.3^AG/AG^ IHCs. (**A**) Single confocal plane of a representative Ca_V_1.3^WT/WT^ IHC showing TAMRA conjugated dimeric Ribeye/Ctbp2 peptide and Fluo4-FF fluorescence. Black and grey colors of voltage ramp stimuli and corresponding whole cell Ca^2+^ currents show IHC response to two voltage ramp depolarizations. Intensity-time profiles of single Ca^2+^ hotspots from one IHC are shown with different colors. (**B**) Average fluorescence-voltage relationships of Ca^2+^ influx at single AZs of Ca_V_1.3^WT/WT^ and Ca_V_1.3^AG/AG^ IHCs. Shaded areas show ± SEM. (**Bi**) The maximal Ca^2+^ influx amplitude (ΔF/F_0 max_) at single AZs is reduced in IHCs of Ca_V_1.3^AG/AG^ mice. (**C**) Fractional activation curves of Ca^2+^ channels at individual AZs. Thick, dark lines show the averages, lighter colors represent individual curves. (**Ci**) The voltage sensitivity of Ca^2+^ channel activation at individual AZs is increased in Ca_V_1.3^AG/AG^ IHCs. (**Cii**) Voltage of half maximal activation (V_half_) of Ca^2+^ channels calculated from the Boltzmann fits in (C) is hyperpolarized in Ca_V_1.3^AG/AG^ IHCs. (**D**) The spatial gradient of maximal Ca^2+^ influx is collapsed in IHCs of Ca_V_1.3^AG/AG^ IHCs. (**E**) The spatial gradient of voltage of half maximal activation is maintained in IHCs of Ca_V_1.3^AG/AG^ IHCs. Polar plots in (D and E) show the positions of individual AZs in Ca_v_1.3^WT/WT^ (left) and Ca_V_1.3^AG/AG^ (right) IHCs. Pseudo color scales represent maximal Ca^2+^ influx amplitude (D) and V_half_ (E). Box plots compare ΔF/F_0 max_ (D) and V_half_ (E) at pillar and modiolar AZs in IHCs of Ca_V_1.3^WT/WT^ and Ca_V_1.3^AG/AG^ mice. Data were acquired from N = 8 (Ca_V_1.3^WT/WT^), 6 (Ca_V_1.3^AG/AG^) mice. Box-Whisker plots with individual data points overlaid show median, 25^th^ and 75^th^ percentiles (box), 10^th^ and 90^th^ percentiles (whiskers). Statistical significances were determined using two-tailed Wilcoxon rank-sum test for data in (Bi), (Ci), (Cii), (D) and (E). Significances are reported as *p < 0.05, **p < 0.01.

**Fig. 2.**
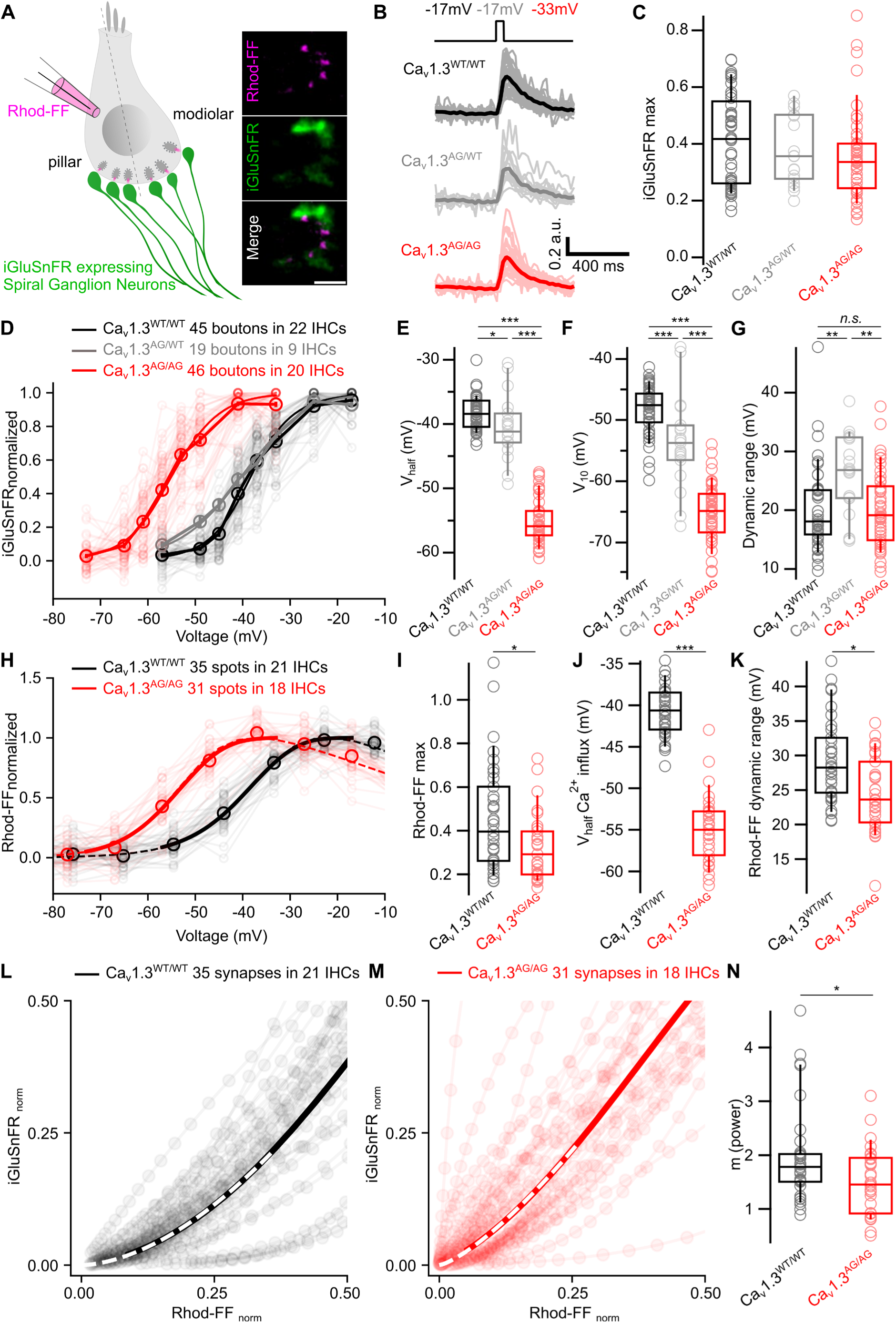
Activation of glutamate release at IHC synapses occurs at lower voltages in Ca_V_1.3^AG/WT^ and Ca_V_1.3^AG/AG^ mice. (**A**) Average projection of ΔF images of Rhod-FF fluorescence from multiple confocal planes of a representative Ca_V_1.3^WT/WT^ IHC (top). Average projection of ΔF images of iGluSnFR fluorescence from 50 ms stimulation pulses of multiple voltages at a single IHC plane (middle). Scale bar = 5 μm. (**B**) Average ΔF/F_0_ traces of iGluSnFR fluorescence. Individual traces are shown in lighter colors. (**C**) Maximal amplitude of iGluSnFR fluorescence (ΔF/F_0 max_) is not affected at Ca_V_1.3^AG/WT^ and Ca_V_1.3^AG/AG^ boutons. (**D**) Average voltage dependence of normalized iGluSnFR signal at single boutons fitted with Boltzmann function. Data from individual boutons are shown only for Ca_V_1.3^WT/WT^ and Ca_V_1.3^AG/AG^ animals. (**E** and **F**) V_half_ (E) and V_10_ (F) of the glutamate release show a gene-dose dependent hyperpolarized shift in Ca_V_1.3^AG/WT^ and Ca_V_1.3^AG/AG^ IHCs compared to Ca_V_1.3^WT/WT^ IHCs. (**G**) The dynamic range of single synaptic glutamate release is increased in Ca_V_1.3^AG/WT^ IHCs. (**H**) Voltage dependence of normalized Ca^2+^ influx at single synapses. Individual traces are shown with lighter colors. Dotted lines show the average modified Boltzmann function fit. Solid, thick lines show the range of the fits matching to the voltage dependency range of iGluSnFR recordings. (**I**) The Rhod-FF ΔF/F_0 max_ is reduced at AZs of Ca_V_1.3^AG/AG^ IHCs. (**J**) V_half_ of Ca^2+^ influx is hyperpolarized in Ca_V_1.3^AG/AG^ IHCs. (**K**) The dynamic range of Ca^2+^ influx at single AZs is reduced in Ca_V_1.3^AG/AG^ IHCs. (**L** and **M**) Synaptic transfer function at single synapses in Ca_V_1.3^WT/WT^ (L) and Ca_V_1.3^AG/AG^ (M) IHCs. Thick, solid line shows the average. White, dotted line represents the power function fitted to the first 25% of the average glutamate release. **n,** The power (*m*) of the Ca^2+^ signal – glutamate release relationship (apparent Ca^2+^ dependence of release) is reduced at the AZs of Ca_V_1.3^AG/AG^ IHC. Data were acquired from N = 12 (Ca_V_1.3^WT/WT^), 5 (Ca_V_1.3^AG/WT^), 9 (Ca_V_1.3^AG/AG^) mice. Box-Whisker plots with individual data points overlaid show median, 25^th^ and 75^th^ percentiles (box), 10^th^ and 90^th^ percentiles (whiskers). Statistical significances were determined using Kruskal-Wallis test for (C), one-way ANOVA followed by Tukey’s HSD for (E) and (F), Kruskal-Wallis followed by Dunn’s test for (G), two-tailed Wilcoxon rank-sum test for (I) and (N), two-tailed t-test for (J) and (K). Significances are reported as *p < 0.05, **p < 0.01, ***p < 0.001.

In order to assess the consequences of reduced number and hyperpolarized activation of Ca^2+^ channels for glutamate release at IHC AZs, we expressed the glutamate sensor iGluSnFR (*39*) in SGN terminals (*4*) (Fig. 2A). The iGluSnFR signal evoked by strong depolarizations (50 ms to -17 mV for Ca_V_1.3^WT/WT^/Ca_V_1.3^AG/WT^ and -33 mV for Ca_V_1.3^AG/AG^) of Ca_V_1.3^AG/AG^ synapses (iGluSnFR_max_) tended to be lower than that of Ca_V_1.3^WT/WT^ synapses without reaching statistical significance (Ca_V_1.3^AG/WT^ fell in-between) (Fig. 2B and C). In contrast to this, but consistent with results obtained with Fluo4-FF dye, presynaptic Ca^2+^ influx of Ca_V_1.3^AG/AG^ synapses reported with the red Ca^2+^ indicator Rhod-FF (k_D_: 320 µM) was significantly reduced (Fig. 2I). The operating range of glutamate release was shifted to lower voltages in Ca_V_1.3^AG/AG^ synapses (-18 mV shift of V_half_, Fig. 2E). We detected significant iGluSnFR signals in Ca_V_1.3^AG/AG^ mice with depolarizations as weak as -65 mV (V_10%_, Fig. 2D and F) compared to -53 mV for Ca_V_1.3^AG/WT^ and -47 mV for Ca_V_1.3^WT/WT^ mice. Interestingly, the activation curves of Ca_V_1.3^AG/WT^ and Ca_V_1.3^WT/WT^ synapses differed only for depolarizations to less than -40 mV, which might result from differences of the Ca_V_1.3^AG/WT^ synapses in expressing WT and AG channels. This could also explain the greater dynamic range of the transfer function found with Ca_V_1.3^AG/WT^ synapses compared to the other genotypes (Fig. 2G). Indeed, by summing fractions of whole cell Ca^2+^ channel activation curves (fig. S1C) from Ca_V_1.3^WT/WT^ and Ca_V_1.3^AG/AG^ IHCs we found that combining 83% WT channels and only 17% AG channels best explained voltage-dependent activation of Ca^2+^ currents in Ca_V_1.3^AG/WT^ IHCs. We found the voltage-dependence of the iGluSnFR signal to closely follow that of Ca^2+^ channel activation for Ca_V_1.3^WT/WT^ synapses (Fig. 2D-G, H, J, K). Next, we related the iGluSnFR signals obtained for different levels of IHC depolarization to the corresponding Ca^2+^ signals to characterize the apparent Ca^2+^ dependence of glutamate release. We note that this protocol changes Ca^2+^ influx primarily via changing the open probability and to a lesser extent via the single channel current. The lower power (*m ∼* 1.5) compared to that of the intrinsic Ca^2+^ dependence of exocytosis (*m* ∼ 2.5) at synapses of Ca_V_1.3^WT/WT^ mice (*4*) is thought to reflect a tight Ca^2+^ nanodomain-like control of synaptic vesicle (SV) release by one or few Ca^2+^ channels with an effective coupling distance of ∼15 nm (*40*, *41*). Ca^2+^ nanodomain-like control of SV release was maintained with even lower power, hence, tighter coupling of Ca^2+^ influx and release at Ca_V_1.3^AG/AG^ AZs (Fig. 2L-N).

Next, we addressed the impact of shifting the synaptic transfer function of Ca_V_1.3^AG/AG^ synapses to lower voltages on spontaneous and sound-evoked firing of SGNs *in vivo.* We first recorded auditory brainstem responses in response to acoustic clicks (ABR, Fig. 3A). The first wave (wave I) reports the firing of the SGN population and its amplitude reflects both the number and firing synchrony of the activated SGNs. The ABR threshold was significantly lower in Ca_V_1.3^AG/AG^ mice than in Ca_V_1.3^WT/WT^ mice, which is consistent with the lower activation threshold of Ca_V_1.3^AG^ channels (Fig. 3B). Wave I amplitude was mildly increased at near-threshold 40 dB acoustic click stimulation, but was not systematically changed across the other sound pressure levels in Ca_V_1.3^AG/AG^ mice (Fig. 3C). This suggests increased firing rates of SGNs at weak stimulations, possibly limited by the steady-state inactivation of the channels. Ca_V_1.3^AG/AG^ mice did not tolerate the urethane/xylazine anesthesia very well and hence, we turned to isoflurane anesthesia for the more demanding recordings from single SGNs (see Materials and Methods). We note that isoflurane inhibits voltage-gated Ca^2+^ channels (*42*) and cannot exclude that the higher open probability renders Ca_V_1.3^AG^ channels more susceptible to isoflurane. This might artificially raise auditory thresholds and obscure differences between Ca_V_1.3^AG/AG^ and Ca_V_1.3^WT/WT^ mice. Indeed, ABR thresholds and wave I amplitudes of both genotypes were not significantly different from each other under isoflurane (fig. S7A-C). Moreover, SRs were strongly reduced in Ca_V_1.3^WT/WT^ SGNs recorded in isoflurane compared to urethane/xylazine (fig. S7D). Despite all this, SR was increased in Ca_V_1.3^AG/AG^ mice and even in Ca_V_1.3^AG/WT^ mice (Fig. 3D). Sound evoked SGN firing was largely intact: frequency tuning and thresholds (Figs. 3E, F, Fi and fig. S7J), and peak firing rates (Fig. 3G, Hi) were not significantly altered in Ca_V_1.3^AG/AG^ or Ca_V_1.3^AG/WT^ mice. Adapted spike rates in Ca_V_1.3^AG/AG^ SGNs were reduced likely reflecting enhanced inactivation of Ca_V_1.3^AG^ channels (Fig. 3H, Hii), which was further tested by 1.5 s long tone bursts (fig. S7E, F, Fi, Fii). The dynamic range of the rate code was comparable across the genotype with a non-significant trend toward larger ranges for Ca_V_1.3^AG/WT^ SGNs, that would be as expected for AZs mixing Ca_V_1.3^AG^ and Ca_V_1.3^WT^ channels (fig. S7G-I). In summary, at least when analyzed in isoflurane, it appears that the cochlea manages to maintain sound encoding at control conditions, despite a massive change in Ca_V_1.3 gating that manifests itself at the level of spontaneous synaptic transmission and SGN firing around sound threshold. The increased SGN SR – evident even under isoflurane – and reduced ABR threshold (urethane/xylazine) demonstrate the impact of presynaptic Ca^2+^ channel gating in IHCs on spontaneous and sound-evoked SGN firing. The graded hyperpolarized shift of IHC synapse activation and ensuing increase of spontaneous SGN firing demonstrated here for Ca_V_1.3^AG^ in a gene dose dependent manner lends support for the hypothesis that the physiologically observed differences of AZs in Ca_V_1.3 activation contribute to SGN firing diversity. It is intriguing that Ca_V_1.3^AG/AG^ IHCs retained the pillar-modiolar AZ gradient of voltage operating range despite the massive hyperpolarized shift. Hence, while Ca_V_1.3^AG^ IHCs homeostatically and likely cell-autonomously downregulate the abundance of Ca_V_1.3 at AZs, the position-dependent regulation of the voltage-dependence of Ca_V_1.3 activation does not seem to target absolute values and might involve cell-non-autonomous signaling.

**Fig. 3.**
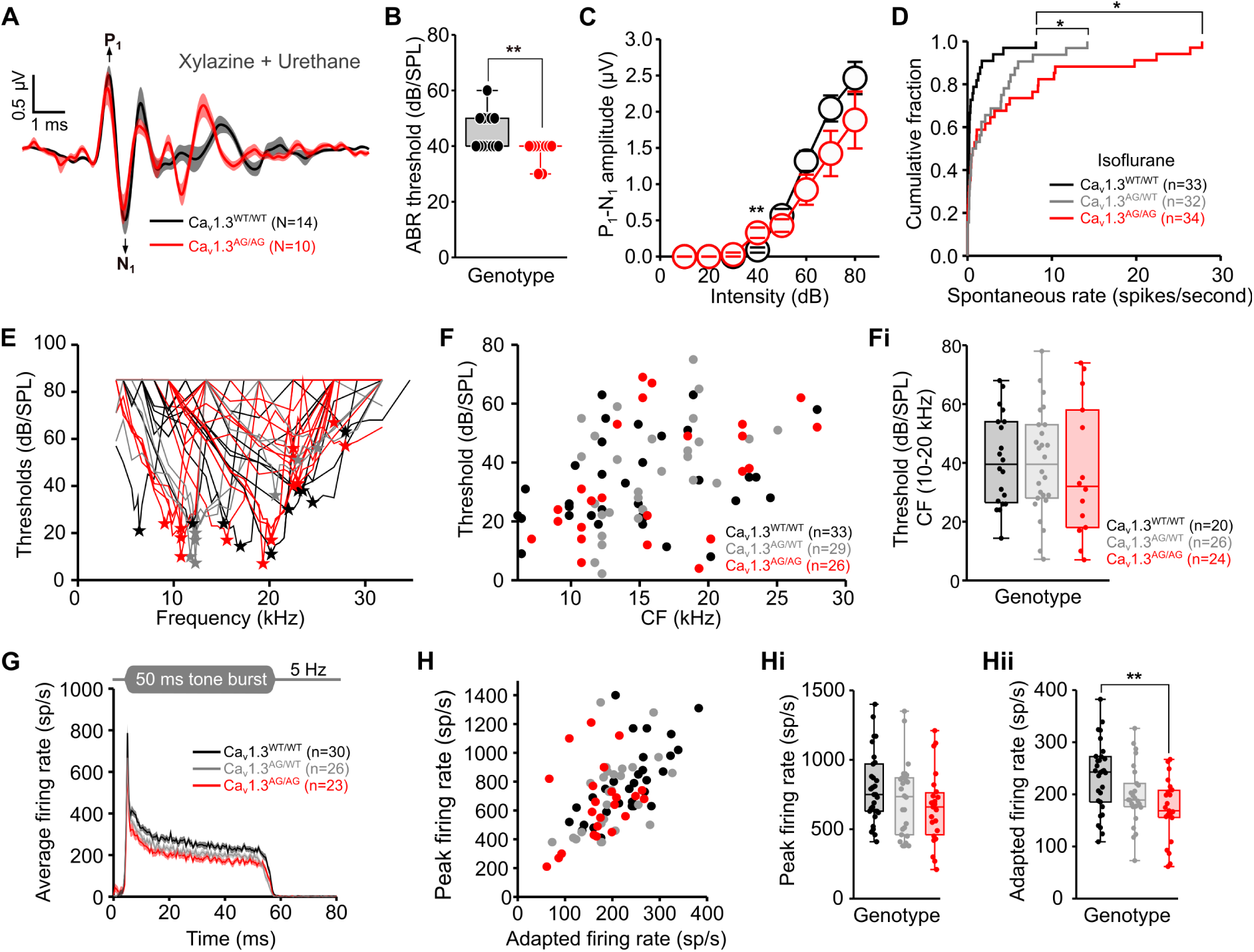
Increased spontaneous rates in SGNs of Ca_V_1.3^AG/WT^ and Ca_V_1.3^AG/AG^ mice. (**A**) Average ABR waveforms in response to 80 dB clicks recorded in mice under xylazine/urethane anesthesia. Shaded areas show ± SEM. (**B**) ABR thresholds in response to click stimuli are lower in Ca_V_1.3^AG/AG^ mice compared to Ca_V_1.3^WT/WT^ littermates. (**C**) ABR P_1_-N_1_ amplitude was significantly bigger in Ca_V_1.3^AG/AG^ at near threshold stimulation (40 dB [peak equivalent]) but similar to Ca_V_1.3^WT/WT^ across other sound pressure levels. (**D**) SRs of SGNs recorded from mice under isoflurane anesthesia show a relative increase of SRs in Ca_V_1.3^AG/WT^ and Ca_V_1.3^AG/AG^ mice compared to Ca_V_1.3^WT/WT^. (**E**) Frequency tuning curves of all recorded putative SGNs with the characteristic frequency (CF)/best threshold marked by stars. (**F** and **Fi**) Thresholds at CF between 10 and 20 kHz (Fi) are comparable in Ca_V_1.3^WT/WT^, Ca_V_1.3^AG/WT^ and Ca_V_1.3^AG/AG^ mice. (**G**) Average PSTH in response to 50 ms stimulation at the CF, 30 dB above the threshold level and stimulation rate of 5 Hz. Shaded areas show ± SEM. (**H**, **Hi** and **Hii**) Onset firing rates (calculated from PSTH as the bin with the highest rate at the sound onset) are not changed (Hi) but the adapted firing rates (averaged firing rates at 35-40 ms after the sound onset) are decreased in SGNs of Ca_V_1.3^AG/AG^ mice (Hii). Single unit recordings are obtained from N = 6 (Ca_V_1.3^WT/WT^), 3 (Ca_V_1.3^AG/WT^), 6 (Ca_V_1.3^AG/AG^) mice. Box-Whisker plots with individual data points overlaid show median, 25^th^ and 75^th^ percentiles (box), and the range (whiskers). Statistical significances were determined using two-tailed Wilcoxon rank-sum test for (B), two-tailed Wilcoxon rank-sum test for each sound level for (C), Kruskal-Wallis test followed by Tukey-Kramer multiple comparison test for (D) and (Hii), Kruskal-Wallis test for (Fi) and (Hi). Significances are reported as *p < 0.05, **p < 0.01.

Based on our findings in Ca_V_1.3^AG/WT^ mice, we postulate higher acoustic sensitivity or normal hearing in patients affected by the Ca_V_1.3^AG^ mutation. Inspired by the gain of IHC AZ function and increased spontaneous SGN firing in Ca_V_1.3^AG/WT^ mice, we addressed the clinically relevant question if patients might face a higher risk of noise-induced synaptopathy (*31–33*). This is also called hidden hearing loss, as it is not detectable by common clinical hearing tests of acoustic sensitivity, because threshold is determined by the function of high SR fibers that are better maintained upon noise exposure (*32*). We turned to electron and immunofluorescence microscopy (EM and confocal microscopy) to study synapse number and morphology across the cochlear turns in Ca_V_1.3^AG/AG^ mice around the onset of hearing and in 1-2 months old adults, when rearing in quiet (see Materials and Methods) and standard-acoustic environment of the animal facility. When reconstructing IHCs residing at various cochlear segments of normally reared Ca_V_1.3^AG/AG^ and Ca_V_1.3^WT/WT^ mice by serial block-face scanning EM (SBEM, Fig. 4A-C), we found a comparable number of SGN terminals contacting the IHC between two groups (Fig. 4D). However, in the mid- and basocochlear regions only half of terminals were associated with ribbons of Ca_V_1.3^AG/AG^ IHCs (Fig. 4E). In agreement with the immunofluorescence data (fig. S4B) the ribbons of apical IHCs were smaller, whereas the remaining ribbons of mid- and basocochlear IHCs were significantly larger (Fig. 4F). We further assessed the morphology of the ribbons by transmission electron microscopy and found a higher abundance of ribbons with a hollow core (fig. S8). Immunohistochemistry confirmed these observations for 2- and 9-months old mice (fig. S9B, Bi, C, Ci), but also revealed that Ca_V_1.3^AG/AG^ IHCs start out with a normal complement of ribbons at the onset of hearing (fig. S9A and Ai) and maintain them better when quiet-reared in particular in the basal cochlea (fig. S9D and Di). Collectively, this data indicates that Ca_V_1.3^AG/AG^ IHCs partially shed their ribbons even at the ambient noise levels of standard animal husbandry, while a potential excitotoxic damage to SGNs was not evident. We cannot rule that excitotoxic damage to SGNs occurred but was repaired by axonal regrowth and formation of new synapses. However, we consider the partial loss of ribbons more likely to represent a selective presynaptic remodeling that might involve Ca^2+^ dependent mitochondrial signaling to ribbons (Fig. 5B) (*43*, *44*) and goes hand in hand with IHC showing a homeostatic reduction of Ca_V_1.3 channels (figs. S3, S4). SGNs showed an increase in the total mitochondrial volumes in their postsynaptic terminals and peripheral neurites in Ca_V_1.3^AG/AG^ that we consider to reflect increased synaptic transmission and/or SGN activity (fig. S10) (*20*). While we did not find a significant ribbon loss in the Ca_V_1.3^AG/WT^ mouse model of human patients (fig. S11), we caution that their ears could be more susceptible to noise-induced hearing loss (Fig. 5B). Future work will need to test this hypothesis with acute and chronic exposure of the mouse model to different noise levels. Finally, further probing the impact of the Ca_V_1.3^AG^ on sound encoding will benefit from identifying anesthesia tolerated by Ca_V_1.3^AG/AG^ mice that does not affect Ca_V_ channels. In summary, our findings support the hypothesis that the voltage-dependence of Ca_V_ gating at a given IHC-SGN synapse governs spontaneous and sound-evoked SGN firing and indicate that heterogeneity of Ca_V_ gating among the synapses contributes to SGN diversity.

**Fig 4.**
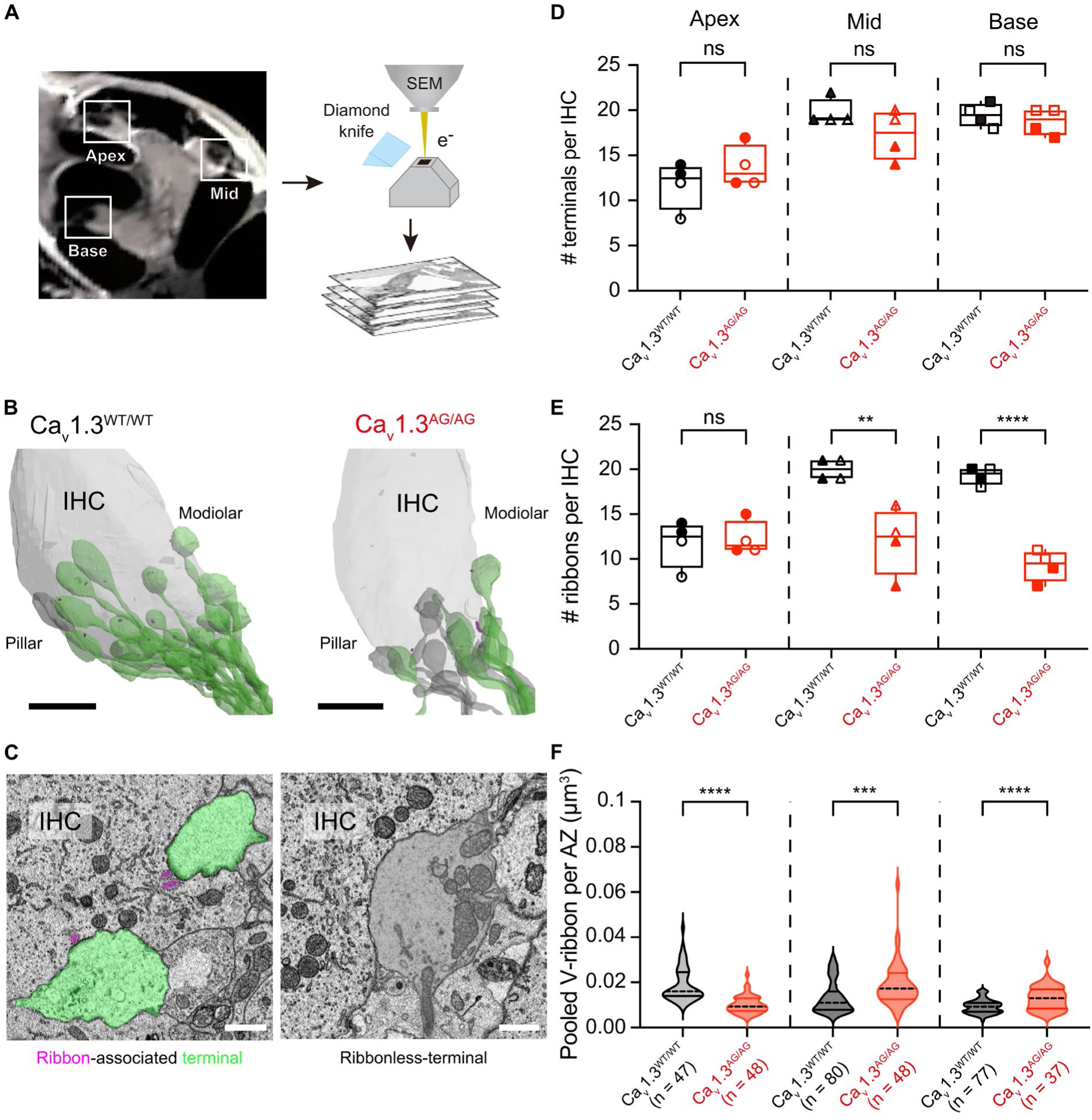
Loss of synaptic ribbons at a subset of IHC AZs in Ca_V_1.3^AG/AG^ mice. (**A**) Schematic illustration of SBEM imaging at apical, middle, and basal segments of the mouse cochlea. (**B**) 3D rendering of reconstructed Ca_V_1.3^WT/WT^ (left) and Ca_V_1.3^AG/AG^ (right) IHCs contacted by SGN terminals with (green) and without (grey, “ribbonless”) associated ribbons (magenta). Scale bar = 5 µm. (**C**) Representative electron micrographs of ribbon-associated (green, left) and ribbonless (grey, right) SGN terminals. Scale bar = 1 µm. (**D**) Average numbers of SGN terminals contacting an IHC are comparable between Ca_V_1.3^AG/AG^ and Ca_V_1.3^WT/WT^ mice. (**E**) Average ribbon counts are significantly smaller in Ca_V_1.3^AG/AG^ IHCs compared to Ca_V_1.3^WT/WT^ at mid- and basal cochlear segments but not in the apex. (**F**) Mean volumes of ribbon are larger in Ca_V_1.3^AG/AG^ IHCs compared to Ca_V_1.3^WT/WT^ IHCs at mid- and basal cochlea segments, whereas apical Ca_V_1.3^AG/AG^ IHCs appear to have exclusively small ribbons. Each tonotopic location of each genotype represents data from N = 2 mice. Box-Whisker plots with individual data points overlaid show median, 25^th^ and 75^th^ percentiles (box), and the range (whiskers). Statistical significances were determined using two-tailed t-test for (D), (E) and (F). Significances are reported as **p < 0.01, ***p < 0.001, ****p < 0.0001.

**Fig 5.**
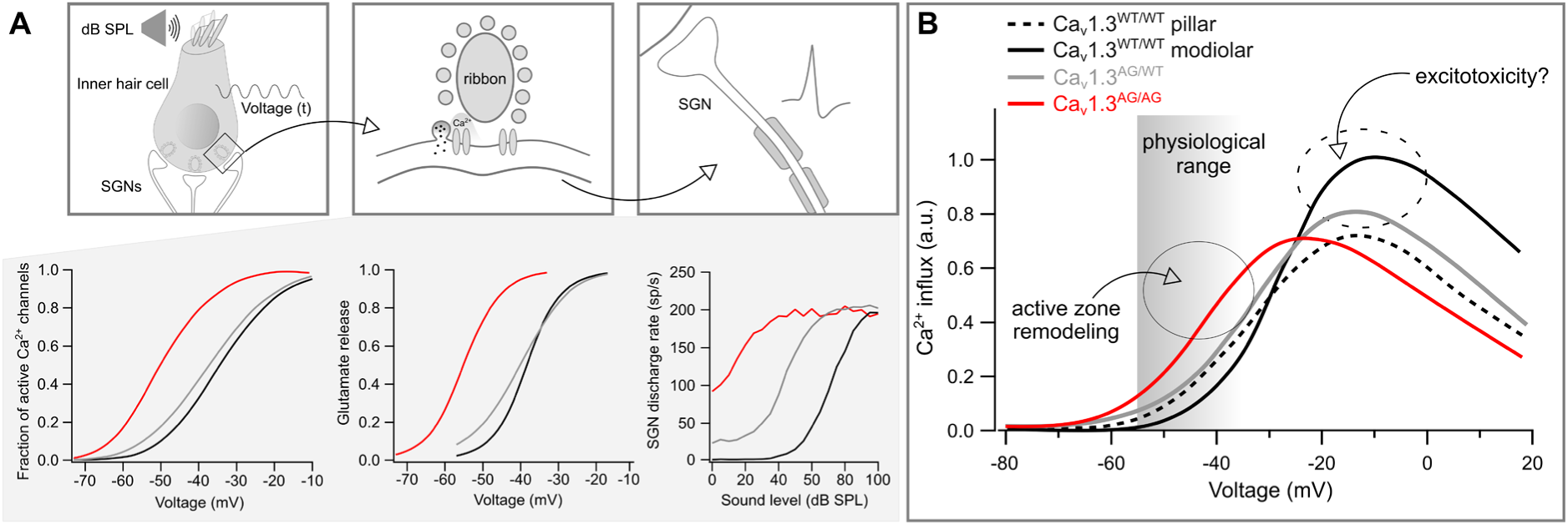
Ca_V_1.3 gating contributes to SGN firing diversity and synaptic vulnerability. (**A**) SGN spontaneous rates and thresholds vary according to the voltage dependence of presynaptic Ca^2+^ channel activation and glutamate release in a gene-dose dependent manner. SGN rate-level functions were modeled according to Meddis et al., 1990 (see Supplementary Information, fig. S11, Table S1). Upper panels summarize the path from sound to SGN code. (**B**) Hyperpolarzed activation of Ca_V_1.3 channels in Ca_V_1.3^AG/AG^ mice leads to Ca^2+^ influx at IHC AZs, exeeding that for both pillar and modiolar AZs in WT IHCs at physiological voltages. This results in homeostatic reduction of Ca^2+^ channels and a general active zone remodeling, but does not reach excitotoxic levels of Ca^2+^ influx and glutamate release at the ambient sound levels of the animal facility. Excitotoxic damage is likely to occur at large receptor potentials during intense sound stimulation and may preferentially affect modiolar synapses that harbor a greater Ca^2+^ channel complement in WT IHCs at non-physiological stimulations, such as moderate noise exposure (shown with the dashed ellipsoid).

## Materials and Methods

### Animals

Ca_V_1.3^AG^ mice have been previously described (*23*). We used homozygous (Ca_V_1.3^AG/AG^), heterozygous (Ca_V_1.3^AG/WT^) mutants and wild-type (Ca_V_1.3^WT/WT^) control mice of either sex for the experiments. Mice were maintained on C57B6/N background. For certain patch-clamp experiments (capacitance measurements, tail current recordings, patch-clamp combined with Fluo4-FF imaging) C57B6J animals were used along with the littermate controls. Ages of the mice varied from 13 days to 9 months depending on the experiment. Animals were housed and raised either in standard husbandry conditions in IVCs or in open cages placed in an isolated quiet environment with low ambient noise levels, particularly lacking the sound of the ventilation system. All the experiments were approved by the local Animal Welfare Committee of the University Medical Center Göttingen and the Max Planck Institute for Multidisciplinary Sciences, as well as the Animal Welfare Office of the state of Lower Saxony, Germany (LAVES, AZ: 19/3134 and 19/3133).

### Patch-clamp

For *ex-vivo* physiology, we dissected 3/4 of the apical turn of the organ of Corti from 2-4-week-old mice. For the experiments, where ruptured patch-clamp was combined with Ca^2+^ imaging using Fluo4-FF dye, IHCs were accessed from the modiolar side in order to preserve the general morphology of the cells for later assignment of modiolar and pillar synapses. For the remaining experiments, IHCs were accessed from the pillar side of the organ. Patch pipettes were made using P-97 Flaming/Brown micropipette puller (Sutter Instruments) and borosilicate glass filaments (GB150-8P, GB150F-8P (Science Products), for perforated and ruptured patch-clamp configurations, respectively). For perforated patch-clamp and capacitance recordings, pipette solution contained (in mM): 130 Cs-gluconate, 10 HEPES, 10 TEA-Cl, 10 4-AP, 1 MgCl_2_, 300 μg/ml amphotericin B, pH 7.3, 290 mOsm. For ruptured patch-clamp combined with imaging pipette solution contained (in mM): 111 Cs-glutamate, 1 MgCl_2_, 1 CaCl_2_, 10 EGTA, 13 TEA-Cl, 20 HEPES, 4 Mg-ATP, 0.3 Na-GTP and 1 L-glutathione, pH 7.3, 290 mOsm. Additionally, Ca^2+^ indicator Fluo-4FF (0.8 mM; Life Technologies) and TAMRA-conjugated Ribeye/Ctbp2 binding peptide (10 mM; Biosyntan) or Rhod-FF (0.8 mM, Biomol) were added to the intracellular solution for live imaging. For ruptured patch-clamp to record the variance of the tail currents pipette solution contained (in mM): 130 Cs-gluconate, 1 MgCl_2_, 10 HEPES, 10 TEA, 0.8 EGTA, 0.4 BAPTA, 10 4-AP, 2 Mg-ATP, 0.3 Na_2_-GTP, 2 mg/ml D-glucose, pH 7.2, 300 mOsm. Perforated patch-clamp was performed in extracellular solution containing (in mM): 107 NaCl, KCl 2.8, 1 MgCl_2_, 10 HEPES, 2 CaCl_2_, 35 TEA, 5 4-AP, 1 CsCl, 2 mg/ml D-glucose, pH 7.2, 300 mOsm. Ruptured patch-clamp combined with imaging was performed in extracellular solution containing (in mM): 2.8 KCl, 105 NaCl, 10 HEPES, 1 CsCl, 1 MgCl_2_, 5 CaCl_2_, 35 TEA-Cl, and 2 mg/ml D-glucose, pH 7.2, 300 mOsm. Ruptured patch-clamp for recording the variance of the tail Ca^2+^ currents was performed in extracellular solution containing (in mM): 90 NaCl, 2.8 KCl, 1 MgCl_2_, 10 HEPES, 10 CaCl_2_, 35 TEA 35, 1 CsCl, 5μM BayK, 2 mg/ml Glucose, 300 mOsm. Data acquisition was done using EPC-10 amplifier (HEKA electronics) controlled by PatchMaster software (HEKA). The holding potential of IHCs was set to -97 mV. All recordings were performed at room temperature (20-25 °C).

### Perforated patch-clamp

Pipettes were coated with Sylgard to minimize the capacitive noise. Capacitance measurements from IHCs were performed using Lindau-Neher technique (*45*) as described previously (*46*). Current-voltage (IV) relationships were recorded by applying 10 ms step depolarizations ranging from -97 mV to 63 mV with 5 mV increments. Recordings were leak corrected using p/n protocol. All voltages were corrected offline for the liquid junction potential (17 mV). Recordings where series resistance (R_s_) exceeded 30 MOhm, leak currents exceeded -50 pA at the holding potential and Ca^2+^ current rundown was more than 25%, were discarded from the analysis. To analyze capacitance changes, traces were averaged 400 ms before and after depolarization (skipping 60 ms of initial segment after depolarization).

### Ruptured patch-clamp

Voltage ramp depolarizations ranging from -97 mV to 53 mV or -97 mV to 63 mV during 150 ms were applied to the cells. Leak correction was done using P/n protocol and liquid junction potential of 17 mV was corrected offline. Recordings were discarded from the analysis if the R_s_ exceeded 14 MOhm during the first 3 minutes after breaking into the cell, leak current exceeded -50 pA at holding potential and Ca^2+^ current rundown was more than 25%. For glutamate imaging, 50 ms step depolarizations to various voltages were applied to the cells in pseudorandom order. The recording and analysis of the variance of Ca^2+^ tail current was performed as described before (*47*).

### Functional imaging

Functional imaging was performed using spinning disc confocal unit (CSU22, Yokogawa) mounted on an upright microscope (Axio Examiner, Zeiss). Spinning disk was set to 2000 rpm. We used 63x, 1.0 NA objective (W Plan-Apochromat, Zeiss) and images were acquired with a sCMOS camera (Andor Neo), with a pixel size of 103 nm. The setup was further equipped with 491 nm (Calypso, Cobolt AB) and 561 nm (Jive, Cobolt AB) lasers. Piezo positioner (Piezosystem) was used to acquire images at different Z planes.

### Fluo4-FF and TAMRA imaging

IHCs were loaded with Fluo4-FF Ca^2+^ dye and TAMRA-conjugated Ribeye/Ctbp2 binding dimeric peptide via the patch pipette. First, the cells were scanned from bottom to top by imaging TAMRA fluorescence with 561 nm laser with 0.5 s exposure time and 0.5 μm step size. This allowed us to obtain cell morphology and visualize synaptic ribbons. Next, we recorded Fluo4-FF fluorescence increase at individual synapses by imaging ribbon containing planes with 491 nm laser at 100 Hz while applying voltage ramp depolarizations to the cell. 2 voltage ramps were applied at each plane, one being 5 ms shifted relative to the other.

### iGluSnFR and Rhod-FF imaging

Genetically encoded glutamate sensor iGluSnFR was targeted to spiral ganglion neurons by postnatal round window injections in p5-7 mice using AAV9 virus where iGluSnFR is expressed under the human synapsin promoter (pAAV9.*hSyn*.iGluSnFR.WPRE.SV40, Addgene or produced in our laboratory), as previously described (*4*). Due to the high density of synapses at the basal planes of IHCs it is difficult to distinguish single postsynaptic boutons from one another. For that reason, we chose to image planes closer to the cell nucleus. Once the plane containing iGluSnFR-expressing postsynaptic boutons was located (central plane), we performed Ca^2+^ imaging at the central plane as well as at 2 planes above and 2 planes below the central plane (step size = 0.5 μm). Rhod-FF fluorescence increase was evoked by 2 identical voltage ramp depolarizations at each of the 5 planes and was imaged using 591 nm laser at 100 Hz. Afterwards, iGluSnFR fluorescence was imaged at the central plane with 491 nm at 50 Hz while stimulating the cell with 50 ms depolarizations of the following voltages in pseudo-random order (in mV): −57, −49, −45, −41, −37, −33, −25, −17 for Ca_V_1.3^WT/WT^ and Ca_V_1.3^AG/WT^ and −73, −65, −61, −57, −53, −49, −41, −33 for Ca_V_1.3^AG/AG^ mice.

### Immunohistochemistry and imaging

Cochleae from animals aged 13 days to 9 months were fixed with 4% FA on ice either for 45 minutes to 1 hour or for 10 minutes whenever Ca_V_1.3 channels were immunolabeled for the purpose of performing STED imaging. Cochleae from 1-month-old mice were fixed in glyoxal solution for 30 minutes on ice followed by 30 minutes at room temperature whenever Ca_V_1.3 channels were stained for analyzing modiolar-pillar gradient of Ca_V_1.3 cluster sizes. Glyoxal fixation has been described before (*48*). Cochleae which were used to count the presynaptic ribbons along the tonotopic axis were further decalcified in EDTA (10%, pH = 8) before dissecting apical, middle and basal turns of the organ of Corti.

The following primary antibodies were used: rabbit anti-Homer1 (1:500, 160 002, Synaptic Systems), mouse anti-Ctbp2 (1:200, 612044, BD Biosciences), rabbit anti-Ca_V_1.3 (1:100, ACC-005, Alomone Labs), mouse anti-Bassoon (1:300, ab82958, Abcam), guinea pig anti-RibeyeA (1:500, 192 104, Synaptic Systems), guinea pig anti-Vglut3 (1:500, 135 204, Synaptic Systems), and chicken anti-calretinin (1:200, 214 106, Synaptic Systems).

The following secondary antibodies were used: Alexa Fluor 488 conjugated anti-guinea pig (1:200, A11073, Thermo Fisher Scientific), Alexa Fluor 488 conjugated anti-rabbit (1:200, A11008, Thermo Fisher Scientific), Alexa Fluor 488 conjugated anti-mouse (1:200, A11001, Thermo Fisher Scientific), Alexa Fluor 568 conjugated anti-chicken (1:200, ab175711, Abcam), Alexa Fluor 568 conjugated anti-guinea pig (1:200, A11075, Thermo Fisher Scientific), Alexa Fluor 647 conjugated anti-rabbit (1:200, ab150079, Abcam), Alexa Fluor 647 conjugated anti-rabbit (1:200, A21244, Thermo Fisher Scientific), STAR 580 conjugated anti-mouse (1:200, Abberior, ST635P-1001-500UG), and STAR 635 conjugated anti-rabbit (1:200, Abberior, ST635P-1002-500UG).

Images were acquired using 100x 1.4 NA oil immersion objective and Abberior Instruments Expert Line STED microscope equipped with 488, 561, and 633 nm lasers and 775 nm STED laser. Z-stacks were acquired in confocal mode, while 2D imaging of the synapses was done in 2D STED mode.

### *In-vivo* recordings

#### Surgical procedure

Mice were anesthetized by intraperitoneal injection of xylazine (5 mg/kg) and urethane (1.32 g/kg) or by inhalation of isoflurane via a face mask (5% vol. in O_2_ for induction, 0.6 – 1.5% vol. in O_2_ for maintenance). Analgesia was achieved using buprenorphine (0.1 mg/kg, repeated every 4 hours) and, in the case of isoflurane anesthesia additional carprofen (5 mg/kg, administered only once at the beginning of the experiment) administered subcutaneously. The animals were maintained at 37 °C throughout the experiment using a custom-made heating pad and placed on a vibration isolation table in a sound-proof chamber (IAC GmbH, Niederkrüchten, Germany). The depth of the anesthesia was regularly monitored by the absence of hind limb withdrawal reflexes and additional anesthetic doses were administered as needed. For the juxtacellular recordings from the auditory nerve, a tracheostomy was performed using a silicon tube to ensure breathing throughout the experiment. In case of the isoflurane anesthesia, the face mask delivering the anesthetic was held in close proximity to the face and tracheal opening of the animals throughout this procedure. The animals were then rapidly positioned in a custom-designed stereotactic head holder and a 3D-printed adaptor was attached to the face mask to efficiently deliver the isoflurane directly to the tracheostomy tube until the end of the experiment. The animals were then positioned in a custom-designed stereotactic head holder, after which the pinnae were removed, the scalp was reflected part of the left occipital bone removed. This procedure then allowed for a partial aspiration of the cerebellum to expose the anterior semicircular canal as a landmark for electrode positioning.

#### Auditory brainstem responses (ABRs)

The stimulus generation, presentation, data acquisition, and offline analysis, was performed using NI System (National Instruments, Austin, TX, USA) and custom-written MATLAB software (The MathWorks, Inc.). The ABRs were recorded by needle electrodes placed subcutaneously near the pinna, on the vertex and on the back near the hind limbs.

The difference potential between vertex and mastoid subdermal needles was amplified 10000 times using a custom-designed amplifier, sampled at a rate of 50 kHz for 20 ms, filtered (300– 3000 Hz) and averaged across 500 presentations. Thresholds were determined by visual inspection as the minimum sound intensity that caused a reproducible response waveform in the recorded traces. The first ABR wave (P_1_-N_1_) was detected manually with a custom-written MATLAB script in which the wave was detected for each trace by the user.

#### Juxtacellular recordings from single putative SGNs

The procedure of juxtacellular recordings from SGNs has been described previously (*49*). Glass microelectrodes (∼ 50 MΩ) were advanced through the posterior end of the anteroventral cochlear nucleus using an LN Mini 55 micro-manipulator (Luigs & Neumann, Germany), aiming toward the internal auditory canal. Acoustic stimulation was provided by an open field Avisoft ScanSpeak Ultrasonic Speaker (Avisoft Bioacoustics, Germany). 50 ms noise bursts served as search stimuli. The spiking responses of isolated sound-responsive neurons were detected and recorded using TDT system III hardware and amplified using ELC-03XS amplifier (NPI Electronic, Tamm, Germany), filtered using a bandpass filter (300-3000 Hz). Offline analysis was performed using waveform-based spike detection by a custom-written MATLAB script. Responses from the central SGN axons were identified and distinguished from cochlear nucleus neurons based on their stereotactic position, noise-burst induced firing, peristimulus time histogram (PSTH), regularity of firing, first spike latency, and spike waveform (*50*). The quality of the spikes was rated subjectively and the recordings with low signal-to-noise ratio were excluded from the analysis.

### Electron microscopy

#### Conventional embedding

Conventional embedding and transmission electron microscopy was performed as described before (*51*, *52*). Briefly, the cochleae of 1-month-old Ca_V_1.3^WT/WT^ and Ca_V_1.3^AG/AG^ mice (2 animals per genotype) were fixed for 1 hour on ice with 4% paraformaldehyde and 0.5% glutaraldehyde in PBS (pH 7.4). This was followed by the dissection of the apical turns of the organs of Corti and an additional overnight fixation with 2% glutaraldehyde in 0.1 M sodium cacodylate buffer (pH 7.2) at 4°C. Afterwards, the samples were washed in 0.1 M sodium cacodylate buffer followed by 1% osmium tetroxide treatment (v/v in 0.1 M sodium cacodylate buffer) for 1 hour. After an additional sodium cacodylate and distilled washing step, the samples were placed in 1% uranyl acetate for 1 hour for *en bloc* staining. Subsequently, the samples were washed 3 times in distilled water and dehydrated in a series of ascending concentrations of EtOH. Finally, the samples were embedded in epoxy resin (Agar 100 kit, Plano, Germany) and polymerized for 48 hours at 70°C. 70-75 nm ultrathin sections were obtained from the polymerized blocks using 35° diamond knife (Diatome AG, Biel, Switzerland) and EM UC7 ultramicrotome (Leica Microsystems, Wetzlar, Germany). 2D electron micrographs were acquired at 80 kV using a JEM1011 transmission electron microscope (JEOL, Freising, Germany) equipped with a Gatan Orius SC1000 camera (Gatan, Munich, Germany).

#### Sample preparation for SBEM

For the SBEM experiment, cochlea samples were harvested from four Ca_V_1.3^WT/WT^ (M1, M2, M3, M4) and two Ca_V_1.3^AG/AG^ (M5, M6) mice at the ages of 36-68 postnatal days. The samples were chemically fixed and *en bloc* stained as previously described (*8*).

In short, the animals were decapitated after CO_2_ inhalation under anesthesia. The dissected cochleae were immediately perfused with an ice-cold fixative mixture through the round and oval windows using an infusion pump (Micro4, WPI, Germany). The fixative solution was freshly prepared and contained 4% paraformaldehyde (Sigma-Aldrich, Germany) and 2.5% glutaraldehyde (Sigma-Aldrich, Germany) buffered with 0.08 M cacodylate (pH 7.4, Sigma-Aldrich, Germany). The samples were immersed in the fixative at 4°C for 5 hours and then transferred to a decalcifying solution made of the same fixative and 5% ethylenediaminetetraacetic acid (EDTA, Serva, Germany) for another 5-hour incubation at 4°C. The samples were then washed twice with 0.15 M cacodylate (pH 7.4) for 30 min each, sequentially immersed in 2% OsO_4_ (Sigma-Aldrich, Germany), 2.5% ferrocyanide (Sigma-Aldrich, Germany), and 2% OsO_4_ at room temperature (RT) for 2, 2, and 1.5 hours. After being washed in 0.15 M cacodylate and distilled water (Sartorius, Germany) for 30 min each, the samples were sequentially incubated in filtered 1% thiocarbohydrazide (TCH, Sigma-Aldrich, Germany) solution and 2% OsO_4_ at RT for 1 and 2 hours, as well as in lead aspartate solution (0.03 M, pH 5.0, adjusted with KOH) at 50°C for 2 hours with intermediate two washing steps with distilled water at RT for 30 min each. The sample embedding was initiated with dehydrated through graded pre-cooled acetone (Carl Roth, Germany) series (50%, 75%, 90%, for 30 min each, all cooled at 4°C) and then pure acetone at RT (three times for 30 min each), followed by resin infiltration with 1:1 and 1:2 mixtures of acetone and Spurr’s resin monomer (4.1 g ERL 4221, 0.95 g DER 736, 5.9 g NSA and 1% DMAE; Sigma-Aldrich, Germany) at RT for 6 and 12 hours on a rotator. After being incubated in pure resin for 12 hours, the samples were placed in an embedding mold (Polyscience, Germany) and hardened in a pre-warmed oven at 70°C for 72 hours.

#### Sample trimming and SBEM imaging

The sample blocks were mounted upright along the conical center axis on aluminum metal rivets (3VMRS12, Gatan, UK) and trimmed coronally towards the modiolus using a diamond trimmer (TRIM2, Leica, Germany). According to anatomical landmarks, block faces of about ∼ 600 × 800 mm^2^ with fields of view at the target segments were created using an ultramicrotome (UC7, Leica, Germany). Sample coating of a 30-nm-thick gold layer was done using a sputter coater (ACE600, Leica, Germany). The serial images were acquired using a field-emission scanning EM (Gemini300, Carl Zeiss, Germany) equipped with an in-chamber ultramicrotome (3ViewXP, Gatan, UK) and back-scattered electron detector (Onpoint, Gatan, UK). Focal charge compensation was set to 100% with a high vacuum chamber pressure of 2.8 10^3^ mbar. Nine datasets (M1/a, M2/a, M3/m, M4/m, M1/b, M2/b, M5/m, M6/m, and M5/b) were imaged at 12 nm pixel size and three datasets (M5/a, M6/a, and M6/b) were at 11 nm pixel size for imaging. All datasets were acquired at 50 nm cutting thickness, 2 keV incident beam energy, and 1.5 ms pixel dwell time.

The Ca_V_1.3^WT/WT^ datasets comprised six image stacks, among which two were from the apical cochlear region (M1/a: 2377 slices with each of 9000 × 15000 pixels, M2/a: 2048 slices of 6000 × 6000 pixels), two from mid-cochlea (M3/m: 3072 slices of 9000 × 14000 pixels, M4/m: 2503 slices of 9000 × 15000 pixels), as well as two from basal cochlear region (M2/b: 3072 slices of 6000 × 6000 pixels, M1/b: 3080 slices of 6000 × 9000 pixels). The six stacks of Ca_V_1.3^AG/AG^ datasets contained 1024 slices (7000 × 9000 pixels, apex, M5/a), 2048 slices (5000 × 9000 pixels, apex, M6/a), 2864 slices (9000 × 15000 pixels, mid, M5/m), 2048 slices (10000 × 14000 pixels, mid, M6/m), 3072 slices (7000 × 9000 pixels, base, M5/b), and 2699 slices (7000 × 9000 pixels, base, M6/b). All datasets were aligned along the z-direction using a self-written MATLAB script based on cross-correlation maximum between consecutive slices (*53*) before being uploaded to webKnossos for data visualization and annotation.

#### Ribbon size measurement, synapse counting and mitochondrial analysis

338 ribbon-type synapses were manually annotated in 24 intact IHCs captured by SBEM using webKnossos. The electron-dense region of individual ribbon synapses was manually contoured, and the associated voxels were counted for ribbon volume measurement. In the case of multi-ribbon synapses, all ribbon bodies at a single active zone were summed up to yield the ribbon volume. 67 large vesicle-free boutons were found to contact the IHC basal lateral poles, which were further identified as non-synaptic terminals of SGN based on their characteristic neurite morphology (*54*). Mitochondrial analysis was performed as previously described (*54*).

### Data analysis

#### Ca^2+^ imaging with Fluo4-FF

Images were analyzed using Igor Pro Software (Wavemetrics). Ca^2+^ hotspots were identified by subtracting the average signal of several baseline frames from the average signal of 5 frames during stimulation (ΔF image). The intensities of the 3×3 matrix surrounding the central pixel of the hotspot were averaged across all time points to obtain the intensity profiles of Ca^2+^ influx over time. Afterwards, the background signal, calculated as an average of approximately 60×60 pixel intensities outside the cell was subtracted from the intensity-time profiles and ΔF/F_0_ traces were calculated. The two ΔF/F_0_ traces (one shifted by 5 ms over the other) were combined, plotted against the corresponding voltages (FV curves) and fitted with a modified Boltzmann function. Fractional activation curves were calculated by fitting the linear decay of the fluorescence signal from the FV curves with a linear function (G_max_), dividing the FV fit by the G_max_ line and fitting the resulting curves with Boltzmann function. Maximal Ca^2+^ influx (ΔF/F_0 max_) was calculated by averaging 5 points during the stimulation. The coordinates of the ribbons obtained from the fluorescence of TAMRA conjugated Ctbp2 peptide were transferred from Cartesian coordinate system to cylindrical coordinate system to assign the pillar and modiolar coordinates of the synapses, as previously described (*2*). Data were not considered for the position dependent analysis of the AZs, whenever the morphology of the IHC was deformed.

#### iGluSnFR and Rhod-FF imaging

Images were analyzed using Python software as described before (*4*). Briefly, ΔF image for all depolarizations were calculated by subtracting the average of 10 baseline images right before the stimulation from the average of 5 frames during the stimulation. To detect the regions of interest (ROIs), average projection of ΔF images from all depolarizations was median filtered (filter level 0.5-4) and maximum entropy thresholding was applied. Touching boutons were separated by watershed segmentation. To ensure that the signal at the ROIs originated from a single AZ, only regions which had a corresponding single Ca^2+^ hotspot, or those adequately spaced apart from each other, were analyzed further. Average of all pixels of each ROI was calculated at all time points. Afterwards, the background signal (average of approximately 60×60 pixel intensities outside the cell) was subtracted from the intensity-time profiles and ΔF/F_0_ traces were calculated. Peak detection was performed by smoothing the ΔF/F_0_ traces using Hanning window function (window size 7). For the area under the curve (AUC) calculation, initial segment of ΔF/F_0_ traces before stimulation was fitted with exponential function and the resulting fit was subtracted from ΔF/F_0_ to accommodate for photobleaching. Subsequently, the AUC between stimulation and 20 frames post-stimulation was computed. AUCs of different depolarizations were plotted against voltages, normalized and fitted with Boltzmann function to obtain the threshold of glutamate release defined as 10% of maximal release (V_10_), voltage of half maximal glutamate release (V_half_) and dynamic range of release, defined as the voltage range between 10% and 90% of maximal release.

ΔF images of Rhod-FF signal were calculated by subtracting average of baseline frames from the average of 5 frames during the highest stimulation at each of the 5 planes. ΔF images of 5 planes were averaged to visualize the hotspots. ROI detection was done similar to iGluSnFR ROI detection. Average of all pixels of each ROI was calculated at all time points for all 5 planes. The plane with highest signal intensity was used to calculate ΔF/F_0._ Band-stop filter was applied to ΔF/F_0_ trances to remove 33 Hz noise caused by the spinning disk. 2 ΔF/F_0_ traces corresponding to 2 identical voltage ramp depolarizations were averaged and plotted against voltage. Modified Boltzmann function was fitted to normalized FV curves similar to the ones obtained from Fluo4-FF imaging. Voltage of half maximal Ca^2+^ influx (V_half_) and dynamic range, defined as the voltage range between 10% and 90% of maximal Ca^2+^ influx, were calculated from modified Boltzmann fits. Maximal Ca^2+^ influx (ΔF/F_0 max_) was calculated by averaging 3 points during the stimulation. Ca^2+^ influx-release coupling at IHC single synapses was estimated by relating the Boltzmann fits of normalized glutamate release to the normalized Ca^2+^ influx of matching voltage ranges and fitting the initial 25% of the glutamate release with power function. Data were discarded from further analysis if the fluorescence signal of the individual synapses could not be well separated from one another and the voltage dependence of iGluSnFR and Rhod-FF fluorescence could not be reliably fitted with Boltzmann function (goodness of fit < 0.7).

#### Immunofluorescence analysis

The Ctbp2 and Ca_V_1.3 immunofluorescent puncta in confocal stacks were analyzed using Imaris software (version 9.6.0, Bitplane) automatic spot detection algorithm. The intensities of the ribbons and Ca_V_1.3 clusters were calculated by summating pixel intensities of the 7×7×5 region around the center of mass of each immunofluorescent spot. Spatial gradients of the ribbon and Ca_V_1.3 cluster sizes were analyzed using Imaris and custom written MATLAB scripts, as described before (*21*, *55*, *56*). Data were excluded from the position dependent analysis, whenever the morphology of the IHCs was deformed.

2D STED images of Ca_V_1.3 line-like clusters were analyzed using Igor Pro software. Briefly, 2D Gaussian function was fitted to the clusters to obtain full with at half maxima (FWHM) of long and short axes using genetic fit algorithm (*57*). The brightness and contrast of the representative images were adjusted for visualization using Fiji software.

#### Statistical analysis

Data were analyzed using Igor Pro (Wavemetrics), Python and MATLAB software. For 2 sample comparisons, data were tested for normality and equality of variances using Jarque-Bera and F-test, respectively. Afterwards, two-tailed t-test or Wilcoxon rank-sum test were performed. The latter was used when normality and/or equality of variances were not met. For multiple comparisons 1 way ANOVA followed by Tukey’s Honestly Significant Difference (HSD) multiple comparison test (for normally distributed data) or Kruskal-Wallis followed by Dunn’s multiple comaprison test (for non-normally distributed data) were used. P-values were corrected for multiple comparisons using Holm-Šídák or Bonferroni-Holm method. Data is presented as mean ± standard error of the mean (S.E.M.), unless otherwise stated. Number of animals is reported as N.

## Acknowledgments

We thank Dr. Fangfang Wang for the SBEM data acquisition. We thank Sina Langer, Christiane Senger-Freitag, and Sandra Gerke for expert technical support. We thank Drs. Barbara Vona and Erwin Neher for comments on the MS. We further thank Dr. Bettina Wolf for the assistance with the anesthesia protocol, Dr. Ursula Fünfschilling for the useful discussions on quiet rearing of the mice, Dr. Kathrin Kusch for the virus production, Julius Bahr and Sophia Mutschall for assisting with the SBEM embedding. NK is a member of the Hertha Sponer College of the Cluster of Excellence Multiscale Bioimaging (MBExC). TM is a Max-Planck Fellow at the Max-Planck Institute for Multidisciplinary Sciences.

## Funding

Deutsche Forschungsgemeinschaft (DFG, German Research Foundation) via the Collaborative Research Center 889 (project A02, TM)

Leibniz Program (MO896/5, TM)

Cluster of Excellence (EXC2067) Multiscale Bioimaging (MBExC, TM)

European Research Council through the Advanced Grant ‘DynaHear” under the European Union’s Horizon 2020 Research and Innovation program (grant agreement No. 101054467, TM)

Fondation Pour l’Audition (FPA, RD-2020-10, TM)

Austrian Science Fund (FWF, DOI 10.55776/P35722, JS)

Austrian Science Fund (FWF, DOI 10.55776/P35087, NJO)

National Natural Science Foundation of China (82171133, YH)

Industrial Support Fund of Huangpu District in Shanghai (XK2019011, YH)

Innovative Research Team of High-level Local Universities in Shanghai (SHSMU-ZLCX20211700, YH)

## Author contributions

TM, NK, and YH designed the study. NK performed patch-clamp, functional imaging and immunohistochemistry, analyzed data and participated in auditory system physiology work under supervision of TM. AT performed auditory system physiology and analysis under supervision of NS and TM. SM performed transmission electron microscopy and analysis under supervision of CW and TM. HW, YQ, and YH performed volume electron microscopy and analysis. QF performed patch-clamp and analysis under supervision of TM. NO and JS generated and provided the mouse model as well as participated in discussion. TM, NK, and YH prepared the initial draft of the manuscript. All authors contributed, reviewed and approved the final version of the manuscript for submission.

Conceptualization: TM, NK, YH

Methodology: NK, AT, SM, QF, YQ, HW, TM, NS, CW, YH

Investigation: NK, AT, SM, QF, YQ, HW

Visualization: NK, AT, SM, QF, HW, YH

Funding acquisition: TM, YH, JS, NJO

Project administration: TM, YH

Supervision: TM, YH, NS, CW

Writing – original draft: TM, YH, NK

Writing – review & editing: TM, YH, NK, AT, NS, CW, JS, NJO

## Data and materials availability

All data and code used in the analysis will be made available upon request.

## References and Notes

1. T. Moser, N. Karagulyan, J. Neef, L. M. Jaime Tobón, Diversity matters — extending sound intensity coding by inner hair cells via heterogeneous synapses. EMBO J 42, e114587 (2023).

2. T.-L. Ohn, M. A. Rutherford, Z. Jing, S. Jung, C. J. Duque-Afonso, G. Hoch, M. M. Picher, A. Scharinger, N. Strenzke, T. Moser, Hair cells use active zones with different voltage dependence of Ca^2+^ influx to decompose sounds into complementary neural codes. PNAS 113, E4716–E4725 (2016).

3. A. C. Meyer, T. Frank, D. Khimich, G. Hoch, D. Riedel, N. M. Chapochnikov, Y. M. Yarin, B. Harke, S. W. Hell, A. Egner, T. Moser, Tuning of synapse number, structure and function in the cochlea. Nat Neurosci 12, 444–453 (2009).

4. Ö. D. Özçete, T. Moser, A sensory cell diversifies its output by varying Ca^2+^ influx-release coupling among active zones. EMBO J 40, e106010 (2021).

5. A. V. Kantardzhieva, M. C. Liberman, W. F. Sewell, Quantitative analysis of ribbons, vesicles, and cisterns at the cat inner hair cell synapse: correlations with spontaneous rate. J. Comp. Neurol. 521, 3260–3271 (2013).

6. A. Merchan-Perez, M. C. Liberman, Ultrastructural differences among afferent synapses on cochlear hair cells: correlations with spontaneous discharge rate. J. Comp. Neurol 371, 208–221 (1996).

7. S. Michanski, K. Smaluch, A. M. Steyer, R. Chakrabarti, C. Setz, D. Oestreicher, C. Fischer, W. Möbius, T. Moser, C. Vogl, C. Wichmann, Mapping developmental maturation of inner hair cell ribbon synapses in the apical mouse cochlea. PNAS 116, 6415–6424 (2019).

8. Y. Hua, X. Ding, H. Wang, F. Wang, Y. Lu, J. Neef, Y. Gao, T. Moser, H. Wu, Electron Microscopic Reconstruction of Neural Circuitry in the Cochlea. Cell Reports 34, 108551 (2021).

9. L. Grant, E. Yi, E. Glowatzki, Two Modes of Release Shape the Postsynaptic Response at the Inner Hair Cell Ribbon Synapse. J Neurosci 30, 4210–4220 (2010).

10. B. R. Shrestha, C. Chia, L. Wu, S. G. Kujawa, M. C. Liberman, L. V. Goodrich, Sensory Neuron Diversity in the Inner Ear Is Shaped by Activity. Cell 174, 1229–1246.e17 (2018).

11. S. Sun, T. Babola, G. Pregernig, K. S. So, M. Nguyen, S.-S. M. Su, A. T. Palermo, D. E. Bergles, J. C. Burns, U. Müller, Hair Cell Mechanotransduction Regulates Spontaneous Activity and Spiral Ganglion Subtype Specification in the Auditory System. Cell 174, 1247–1263.e15 (2018).

12. C. Petitpré, H. Wu, A. Sharma, A. Tokarska, P. Fontanet, Y. Wang, F. Helmbacher, K. Yackle, G. Silberberg, S. Hadjab, F. Lallemend, Neuronal heterogeneity and stereotyped connectivity in the auditory afferent system. Nat Commun 9, 3691 (2018).

13. C. Li, X. Li, Z. Bi, K. Sugino, G. Wang, T. Zhu, Z. Liu, Comprehensive transcriptome analysis of cochlear spiral ganglion neurons at multiple ages. Elife 9 (2020).

14. N. Y. S. Kiang, T. Watanabe, Thomas, E.C., Clark, L.F., Discharge Patterns of Single Fibers in the Cat’s Auditory Nerve (MIT Press, Cambridge, Massachusetts, 1965).

15. I. M. Winter, D. Robertson, G. K. Yates, Diversity of characteristic frequency rate-intensity functions in guinea pig auditory nerve fibres. Hear. Res. 45, 191–202 (1990).

16. M. B. Sachs, P. J. Abbas, Rate versus level functions for auditory-nerve fibers in cats: tone-burst stimuli. J. Acoust. Soc. Am. 56, 1835–1847 (1974).

17. M. C. Liberman, Auditory-nerve response from cats raised in a low-noise chamber. J. Acoust. Soc. Am 63, 442–455 (1978).

18. C. L. Adamson, M. A. Reid, R. L. Davis, Opposite actions of brain-derived neurotrophic factor and neurotrophin-3 on firing features and ion channel composition of murine spiral ganglion neurons. J. Neurosci 22, 1385–1396 (2002).

19. A. L. Markowitz, R. Kalluri, Gradients in the biophysical properties of neonatal auditory neurons align with synaptic contact position and the intensity coding map of inner hair cells. Elife 9, e55378 (2020).

20. M. C. Liberman, Single-Neuron Labeling in the Cat Auditory Nerve. Science 216, 1239– 1241 (1982).

21. H. E. Sherrill, P. Jean, E. C. Driver, T. R. Sanders, T. S. Fitzgerald, T. Moser, M. W. Kelley, Pou4f1 Defines a Subgroup of Type I Spiral Ganglion Neurons and Is Necessary for Normal Inner Hair Cell Presynaptic Ca2+ Signaling. J. Neurosci. 39, 5284–5298 (2019).

22. C. Siebald, P. F. Y. Vincent, R. T. Bottom, S. Sun, D. O. J. Reijntjes, M. Manca, E. Glowatzki, U. Müller, Molecular signatures define subtypes of auditory afferents with distinct peripheral projection patterns and physiological properties. Proceedings of the National Academy of Sciences 120, e2217033120 (2023).

23. N. J. Ortner, A. Sah, E. Paradiso, J. Shin, S. Stojanovic, N. Hammer, M. Haritonova, N. T. Hofer, A. Marcantoni, L. Guarina, P. Tuluc, T. Theiner, F. Pitterl, K. Ebner, H. Oberacher, E. Carbone, N. Stefanova, F. Ferraguti, N. Singewald, J. Roeper, J. Striessnig, The human channel gating–modifying A749G CACNA1D (Cav1.3) variant induces a neurodevelopmental syndrome–like phenotype in mice. JCI Insight 8, e162100 (2023).

24. A. Pinggera, A. Lieb, B. Benedetti, M. Lampert, S. Monteleone, K. R. Liedl, P. Tuluc, J. Striessnig, CACNA1D de novo mutations in autism spectrum disorders activate cav1.3 l-type calcium channels. Biological Psychiatry, doi: 10.1016/j.biopsych.2014.11.020 (2015).

25. S. M. Baig, A. Koschak, A. Lieb, M. Gebhart, C. Dafinger, G. Nürnberg, A. Ali, I. Ahmad, M. J. Sinnegger-Brauns, N. Brandt, J. Engel, M. E. Mangoni, M. Farooq, H. U. Khan, P. Nürnberg, J. Striessnig, H. J. Bolz, Loss of Ca(v)1.3 (CACNA1D) function in a human channelopathy with bradycardia and congenital deafness. Nat Neurosci 14, 77–84 (2011).

26. N. J. Ortner, T. Kaserer, J. N. Copeland, J. Striessnig, De novo CACAN1D Ca2+ channelopathies: clinical phenotypes and molecular mechanism. Pflugers Arch - Eur J Physiol 472, 755–773 (2020).

27. S. L. Johnson, Membrane properties specialize mammalian inner hair cells for frequency or intensity encoding. eLife 4, e08177 (2015).

28. D. Oestreicher, S. Chepurwar, K. Kusch, V. Rankovic, S. Jung, N. Strenzke, T. Pangrsic, CaBP1 and 2 enable sustained CaV1.3 calcium currents and synaptic transmission in inner hair cells. bioRxiv [Preprint] (2023). 10.1101/2023.10.16.562475.

29. M. M. Picher, A. Gehrt, S. Meese, A. Ivanovic, F. Predoehl, S. Jung, I. Schrauwen, A. G. Dragonetti, R. Colombo, G. V. Camp, N. Strenzke, T. Moser, Ca2+-binding protein 2 inhibits Ca2+-channel inactivation in mouse inner hair cells. PNAS 114, E1717–E1726 (2017).

30. I. Schrauwen, S. Helfmann, A. Inagaki, F. Predoehl, M. A. Tabatabaiefar, M. M. Picher, M. Sommen, C. Z. Seco, J. Oostrik, H. Kremer, A. Dheedene, C. Claes, E. Fransen, M. H. Chaleshtori, P. Coucke, A. Lee, T. Moser, G. Van Camp, A Mutation in CABP2, Expressed in Cochlear Hair Cells, Causes Autosomal-Recessive Hearing Impairment. Am J Hum Genet 91, 636–645 (2012).

31. S. G. Kujawa, M. C. Liberman, Adding insult to injury: cochlear nerve degeneration after “temporary” noise-induced hearing loss. The Journal of Neuroscience 29, 14077 (2009).

32. A. C. Furman, S. G. Kujawa, M. C. Liberman, Noise-induced cochlear neuropathy is selective for fibers with low spontaneous rates. J. Neurophysiol. 110, 577–586 (2013).

33. Y. Lu, J. Liu, B. Li, H. Wang, F. Wang, S. Wang, H. Wu, H. Han, Y. Hua, Spatial patterns of noise-induced inner hair cell ribbon loss in the mouse mid-cochlea. iScience 27, 108825 (2024).

34. P. Hess, J. B. Lansman, R. W. Tsien, Different modes of Ca channel gating behaviour favoured by dihydropyridine Ca agonists and antagonists. Nature 311, 538–544 (1984).

35. W. M. Roberts, R. A. Jacobs, A. J. Hudspeth, Colocalization of ion channels involved in frequency selectivity and synaptic transmission at presynaptic active zones of hair cells. J Neurosci 10, 3664–3684 (1990).

36. A. Brandt, Few CaV1.3 Channels Regulate the Exocytosis of a Synaptic Vesicle at the Hair Cell Ribbon Synapse. Journal of Neuroscience 25, 11577–11585 (2005).

37. T. Frank, D. Khimich, A. Neef, T. Moser, Mechanisms contributing to synaptic Ca2+ signals and their heterogeneity in hair cells. Proceedings of the National Academy of Sciences 106, 4483–4488 (2009).

38. D. Zenisek, N. K. Horst, C. Merrifield, P. Sterling, G. Matthews, Visualizing synaptic ribbons in the living cell. J. Neurosci 24, 9752–9759 (2004).

39. J. S. Marvin, B. G. Borghuis, L. Tian, J. Cichon, M. T. Harnett, J. Akerboom, A. Gordus, S. L. Renninger, T.-W. Chen, C. I. Bargmann, M. B. Orger, E. R. Schreiter, J. B. Demb, W.-B. Gan, S. A. Hires, L. L. Looger, An optimized fluorescent probe for visualizing glutamate neurotransmission. Nat Meth 10, 162–170 (2013).

40. F. K. Wong, A. R. Nath, R. H. C. Chen, S. R. Gardezi, Q. Li, E. F. Stanley, Synaptic vesicle tethering and the CaV2.2 distal C-terminal. Front Cell Neurosci 8 (2014).

41. T. Pangršič, M. Gabrielaitis, S. Michanski, B. Schwaller, F. Wolf, N. Strenzke, T. Moser, EF-hand protein Ca2+ buffers regulate Ca2+ influx and exocytosis in sensory hair cells. PNAS 112, E1028–E1037 (2015).

42. R. E. Study, Isoflurane Inhibits Multiple Voltage-gated Calcium Currents in Hippocampal Pyramidal Neurons. Anesthesiology 81, 104–116 (1994).

43. H.-T. C. Wong, Q. Zhang, A. J. Beirl, R. S. Petralia, Y.-X. Wang, K. Kindt, Synaptic mitochondria regulate hair-cell synapse size and function. Elife 8, e48914 (2019).

44. L. Sheets, K. S. Kindt, T. Nicolson, Presynaptic CaV1.3 Channels Regulate Synaptic Ribbon Size and Are Required for Synaptic Maintenance in Sensory Hair Cells. J Neurosci 32, 17273–17286 (2012).

45. M. Lindau, E. Neher, Patch-clamp techniques for time-resolved capacitance measurements in single cells. Pflügers Archiv European Journal of Physiology 411, 137– 146 (1988).

46. T. Moser, D. Beutner, Kinetics of exocytosis and endocytosis at the cochlear inner hair cell afferent synapse of the mouse. Proc Natl Acad Sci U S A 97, 883–888 (2000).

47. J. Neef, A. Gehrt, A. V. Bulankina, A. C. Meyer, D. Riedel, R. G. Gregg, N. Strenzke, T. Moser, The Ca^2+^ Channel Subunit beta2 Regulates Ca^2+^ Channel Abundance and Function in Inner Hair Cells and Is Required for Hearing. J Neurosci 29, 10730 (2009).

48. K. N. Richter, N. H. Revelo, K. J. Seitz, M. S. Helm, D. Sarkar, R. S. Saleeb, E. D’Este, J. Eberle, E. Wagner, C. Vogl, D. F. Lazaro, F. Richter, J. Coy-Vergara, G. Coceano, E. S. Boyden, R. R. Duncan, S. W. Hell, M. A. Lauterbach, S. E. Lehnart, T. Moser, T. Outeiro, P. Rehling, B. Schwappach, I. Testa, B. Zapiec, S. O. Rizzoli, Glyoxal as an alternative fixative to formaldehyde in immunostaining and super-resolution microscopy. The EMBO Journal, e201695709 (2017).

49. Z. Jing, M. A. Rutherford, H. Takago, T. Frank, A. Fejtova, D. Khimich, T. Moser, N. Strenzke, Disruption of the presynaptic cytomatrix protein bassoon degrades ribbon anchorage, multiquantal release, and sound encoding at the hair cell afferent synapse. J Neurosci 33, 4456–4467 (2013).

50. A. M. Taberner, M. C. Liberman, Response Properties of Single Auditory Nerve Fibers in the Mouse. J Neurophysiol 93, 557–569 (2005).

51. A. B. Wong, M. A. Rutherford, M. Gabrielaitis, T. Pangršič, F. Göttfert, T. Frank, S. Michanski, S. Hell, F. Wolf, C. Wichmann, T. Moser, Developmental refinement of hair cell synapses tightens the coupling of Ca2+ influx to exocytosis. EMBO J 33, 247–264 (2014).

52. P. Jean, D. L. de la Morena, S. Michanski, L. M. J. Tobón, R. Chakrabarti, M. M. Picher, J. Neef, S. Jung, M. Gültas, S. Maxeiner, A. Neef, C. Wichmann, N. Strenzke, C. Grabner, T. Moser, The synaptic ribbon is critical for sound encoding at high rates and with temporal precision. Elife 7, e29275 (2018).

53. Y. Hua, S. Loomba, V. Pawlak, K.-M. Voit, P. Laserstein, K. M. Boergens, D. J. Wallace, J. N. D. Kerr, M. Helmstaedter, Connectomic analysis of thalamus-driven disinhibition in cortical layer 4. Cell Rep 41, 111476 (2022).

54. Y. Lu, Y. Jiang, F. Wang, H. Wu, Y. Hua, Electron Microscopic Mapping of Mitochondrial Morphology in the Cochlear Nerve Fibers. JARO 25, 341–354 (2024).

55. P. Jean, Ö. D. Özçete, B. Tarchini, T. Moser, Intrinsic planar polarity mechanisms influence the position-dependent regulation of synapse properties in inner hair cells. Proc Natl Acad Sci USA 116, 9084–9093 (2019).

56. N. Karagulyan, T. Moser, Synaptic activity is not required for establishing heterogeneity of inner hair cell ribbon synapses. Front. Mol. Neurosci. 16, 1248941 (2023).

57. M. Sanchez del Rio, G. Pareschi, Global optimization and reflectivity data fitting for x-ray multilayer mirrors by means of genetic algorithms. SPIE Proceedings, 88–96 (2001).

58. R. Meddis, M. J. Hewitt, T. M. Shackleton, Implementation details of a computation model of the inner hair-cell auditory-nerve synapse. The Journal of the Acoustical Society of America 87, 1813–1816 (1990).

59. B. N. Buran, N. Strenzke, A. Neef, E. D. Gundelfinger, T. Moser, M. C. Liberman, Onset coding is degraded in auditory nerve fibers from mutant mice lacking synaptic ribbons. J. Neurosci. 30, 7587–7597 (2010).

