## Supplementary Materials for "Gating of hair cell Ca^2+^ channels governs the activity of cochlear neurons"

**The PDF file includes:**

Supplementary Text
Figs. S1 to S12
Table S1
References

### Supplementary Text

#### IHC-SGN synaptic transmission model

IHC-SGN synaptic transmission and SGN spike generation was modeled according to Meddis et al., 1990 (*58*) using Igor Pro software. In brief, the transmitter release rate/fraction [k(t)], is a sigmoidal function directly dependent on the stimulus level [st(t)]: $k(t)=g(st(t) + A) / (st(t) + A + B)$. The parameter g represents the maximal release rate, A controls the baseline release, as well as the threshold of the release and B controls the saturation and the steepness of the curve. Under the assumption of $Ca^{2+}$ nanodomain control of release (Fig. 4), k(t) reflects the stimulus dependence of $Ca^{2+}$ channel activation. The parameters A and B of the release fraction equation are modified in order to shift the sigmoidal function (Table S1, fig. S12). The release fraction determines the synaptic cleft transmitter content (c). Spike generation is scaled by the cleft transmitter content, firing probability scaling factor (h) and further modulated by absolute (0.8 ms) and relative (2 ms) refractory periods (*59*). The relative refractory period is a random number drawn from a monoexponential distribution with an average value of 2 ms. Rate-level functions are obtained by calculating the adapted firing rates from the simulated PSTH (50 ms stimulation duration).

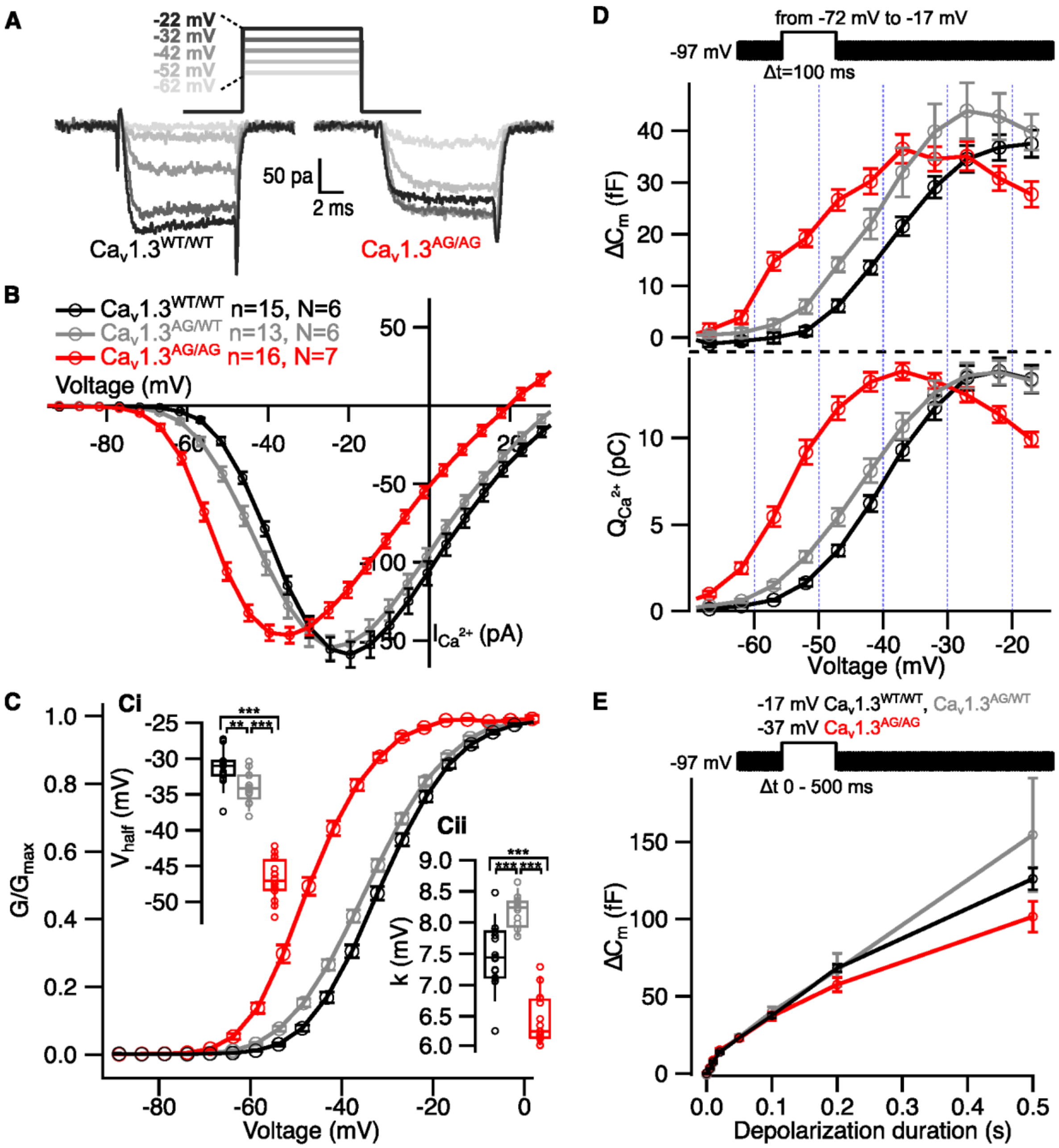


**Fig. S1.**

**Shift to lower voltages and altered voltage sensitivity of $Ca_V1.3$ activation in IHCs of $Ca_V1.3^{AG/WT}$ and $Ca_V1.3^{AG/AG}$ mice.** (**A**) Representative $Ca^{2+}$ current traces from $Ca_V1.3^{WT/WT}$ (bottom left) and $Ca_V1.3^{AG/AG}$ (bottom right) IHCs evoked by step depolarizations (top). (**B**) Whole cell $Ca^{2+}$ current-voltage relationships (IV curves) show comparable maximal $Ca^{2+}$ current amplitude in $Ca_V1.3^{WT/WT}$, $Ca_V1.3^{AG/WT}$, and $Ca_V1.3^{AG/AG}$ IHCs. Error bars show ± SEM (**C**) $Ca^{2+}$ channel activation-voltage relationships calculated from IV-curves show a hyperpolarized shift in $Ca_V1.3^{AG/WT}$ and $Ca_V1.3^{AG/AG}$ IHCs. Error bars show ± SEM. (**Ci**) The voltage of half maximal activation ($V_{half}$) is hyperpolarized in $Ca_V1.3^{AG/WT}$ and $Ca_V1.3^{AG/AG}$ IHCs. (**Cii**) The voltage sensitivity (k) is decreased in $Ca_V1.3^{AG/WT}$ and increased in $Ca_V1.3^{AG/AG}$ IHCs compared to the controls. (**D**) Mean exocytic $\Delta C_m$ and $Ca^{2+}$ current integrals ($Q_{ca}$) evoked by 100 ms pulses of different depolarizations. (**E**) Mean exocytic $\Delta C_m$ in response to different depolarization durations. Data in (B), (C) and (D) is presented as mean ± SEM. Box-Whisker plots with individual data points overlaid show median, 25$^{th}$ and 75$^{th}$ percentiles (box), 10$^{th}$ and 90$^{th}$ percentiles (whiskers). Statistical significances were determined using one-way ANOVA followed by Tukey's Honestly Significant Difference (HSD) for (Ci) and (Cii). Significances are reported as $^{**}p < 0.01$, $^{***}p < 0.001$.

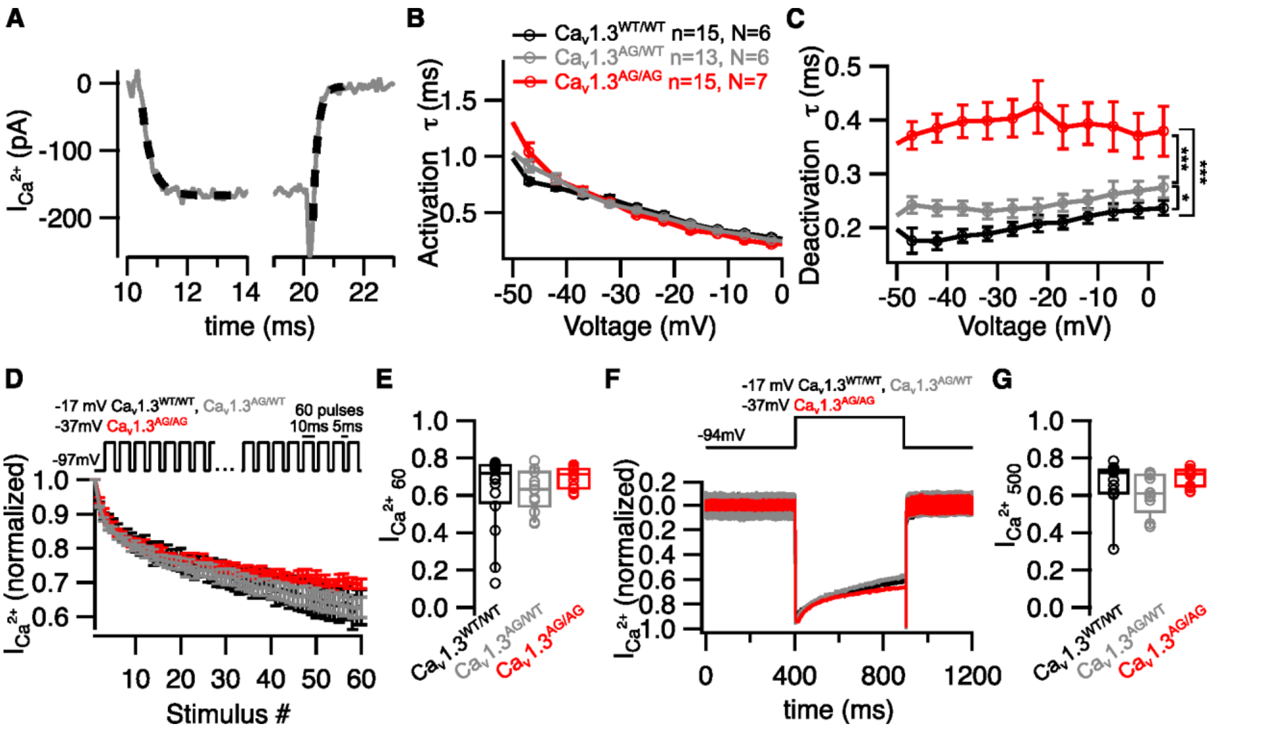


**Fig. S2.**

**Slow deactivation of $Ca^{2+}$ channels in IHCs of Cav1.$3^{AG/WT}$ and Cav1.$3^{AG/AG}$ mice.** (**A**) Activation and deactivation constants of $Ca^{2+}$ currents were obtained by fitting exponential functions (dotted lines) to the first 3 ms of activation and 1 ms of deactivation. (**B**) Cav1.3 activation kinetics across voltages are not changed in IHCs of Cav1.$3^{AG/WT}$ and Cav1.$3^{AG/AG}$ mice. (**C**) The deactivation kinetics are slower in IHCs of Cav1.$3^{AG/WT}$ and Cav1.$3^{AG/AG}$ mice. (**D**) Normalized and averaged Cav1.3 currents measured by applying 60 repetitive stimulations with 10 ms duration and 5 ms interstimulus interval in IHCs of Cav1.$3^{WT/WT}$, Cav1.$3^{AG/WT}$, and Cav1.$3^{AG/AG}$ mice. Error bars show ± SEM. (**E**) The fraction of $Ca^{2+}$ current remaining after 60 repetitive stimulations is comparable in Cav1.$3^{WT/WT}$, Cav1.$3^{AG/WT}$, and Cav1.$3^{AG/AG}$ IHCs. (**F**) Representative normalized $Ca^{2+}$ currents measured by applying 500 ms depolarization at the maximal activation voltage in Cav1.$3^{WT/WT}$, Cav1.$3^{AG/WT}$, and Cav1.$3^{AG/AG}$ IHCs. (**G**) The fraction of $Ca^{2+}$ current remaining after 500 ms depolarization is comparable in Cav1.$3^{WT/WT}$, Cav1.$3^{AG/WT}$, and Cav1.$3^{AG/AG}$ IHCs. Data in (B), (C) and (D) is presented as mean ± SEM. Box-Whisker plots with individual data points overlaid show median, 25th and 75th percentiles (box), 10th and 90th percentiles (whiskers). Statistical significances were determined using one-way ANOVA/Kruskal-Wallis test followed by Tukey's HSD/Dunn's test for each voltage for (B) and (C) and Kruskal-Wallis test for (E) and (G). Significances are reported as *$p < 0.05$, ***$p < 0.001$.

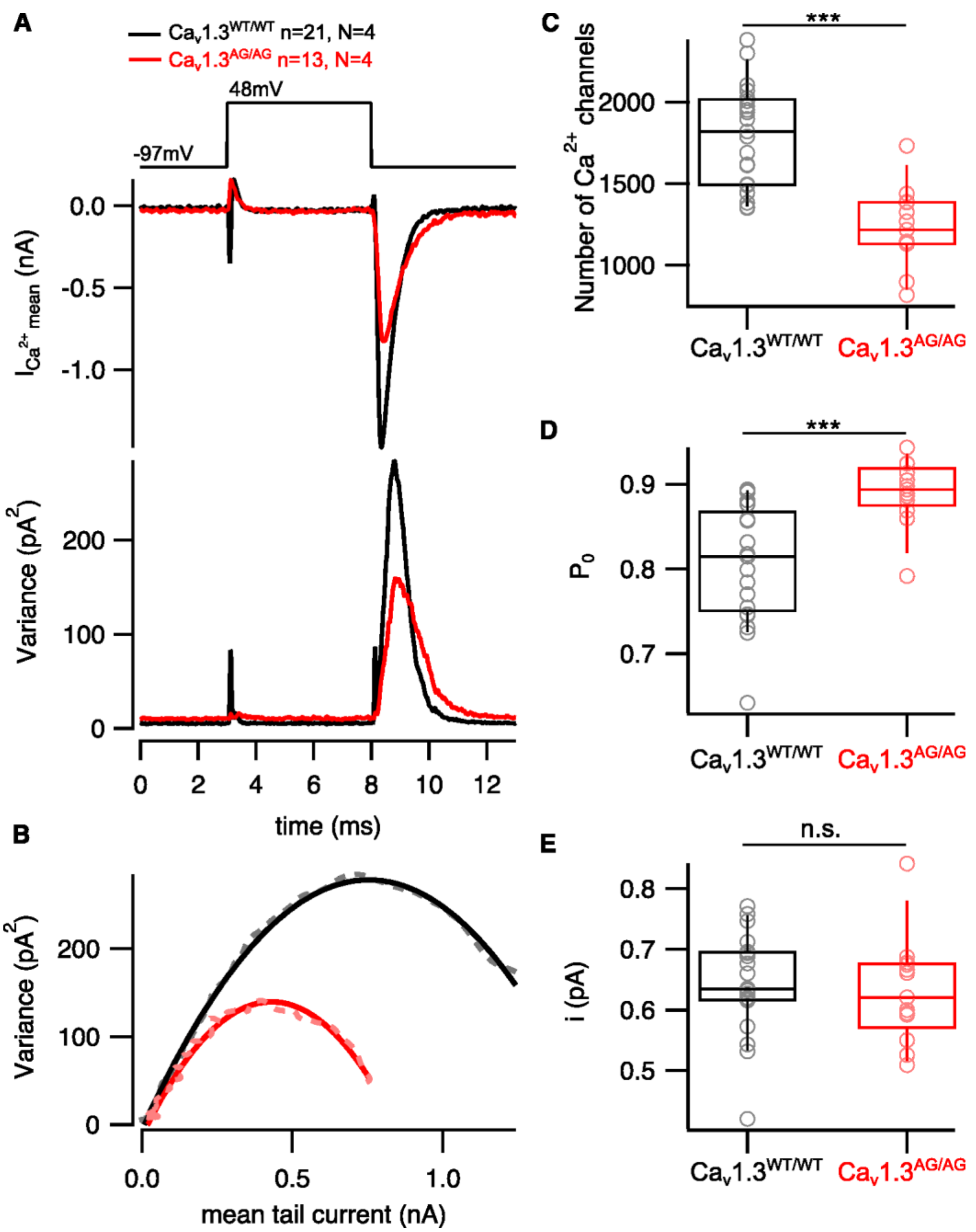


**Fig. S3.**

**Increased open probability and decreased number of Ca$_V$1.3 channels in Ca$_V$1.3$^{AG/AG}$ IHCs.** (**A**) Exemplary mean $Ca^{2+}$ current (middle) evoked by the voltage clamp protocol (top) and variance of the mean current (bottom). Currents were recorded in the presence of 5 µM BayK and 10 mM extracellular $Ca^{2+}$. (**B**) Data from exemplary cells showing variance of the mean $Ca^{2+}$ current plotted against the mean $Ca^{2+}$ current and fitted with a quadratic function. (**C**) The number of activatable $Ca^{2+}$ channels is reduced in Cav1.3$^{AG/AG}$ IHCs. (**D**) $Ca^{2+}$ channels in Cav1.3$^{AG/AG}$ IHCs show higher open probability ($P_o$) compared to Cav1.3$^{WT/WT}$ IHCs. (**E**) Single channel current (i) of $Ca^{2+}$ channels is not changed in IHCs of Cav1.3$^{AG/AG}$ mice. Box-Whisker plots with individual data points overlaid show median, 25$^{th}$ and 75$^{th}$ percentiles (box), 10$^{th}$ and 90$^{th}$

percentiles (whiskers). Statistical significances were determined using two-tailed Wilcoxon rank-sum test for (C), two-tailed t-test for (D) and (E). Significances are reported as ***p < 0.001.

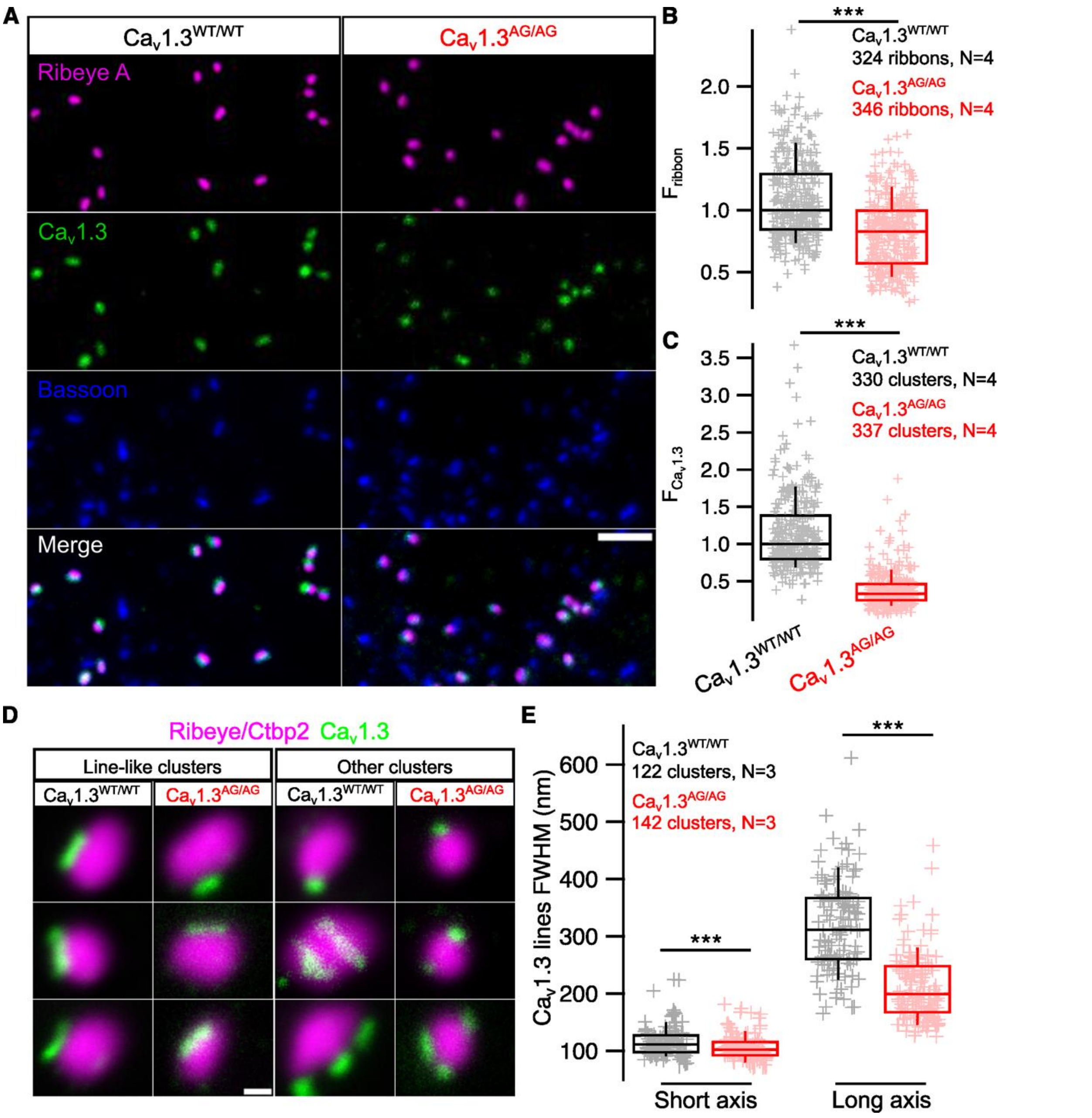


**Fig. S4.**

**Smaller CaV1.3 channel clusters and ribbons at active zones of apical CaV1.3$^{AG/AG}$ IHCs.** (**A**) Maximal intensity projections of confocal stacks acquired from IHC synaptic regions and immunolabeled against Ribeye A, Bassoon and Cav1.3. Scale bar = 2 µm. (**B**) Immunofluorescence intensity of the synaptic ribbons at the apical turn of the cochlea is reduced in Cav1.3$^{AG/AG}$ IHCs. All values were normalized to the median intensity of the Cav1.3$^{WT/WT}$ ribbons. (**C**) Immunofluorescence intensity of synaptic Cav1.3 positive puncta obtained from the confocal stacks is reduced in Cav1.3$^{AG/AG}$ IHCs. All values were normalized to the median intensity of the Cav1.3$^{WT/WT}$ channel clusters. (**D**) Representative images of IHC AZs acquired by STED imaging of AZs immunolabeled for Ribeye/Ctbp2 and Cav1.3. Scale bar = 200 nm. (**E**) Cav1.3 line-like clusters fitted with 2D gaussian function show reduced full with at half maxima

(FWHM) of long and short axes in Cav1.3$^{AG/AG}$ IHCs. Box-Whisker plots with individual data points overlaid show median, 25$^{th}$ and 75$^{th}$ percentiles (box), 10$^{th}$ and 90$^{th}$ percentiles (whiskers). Statistical significances were determined using two-tailed Wilcoxon rank-sum test for data in (B), (C) and (E). Significances are reported as ***$p < 0.001$.

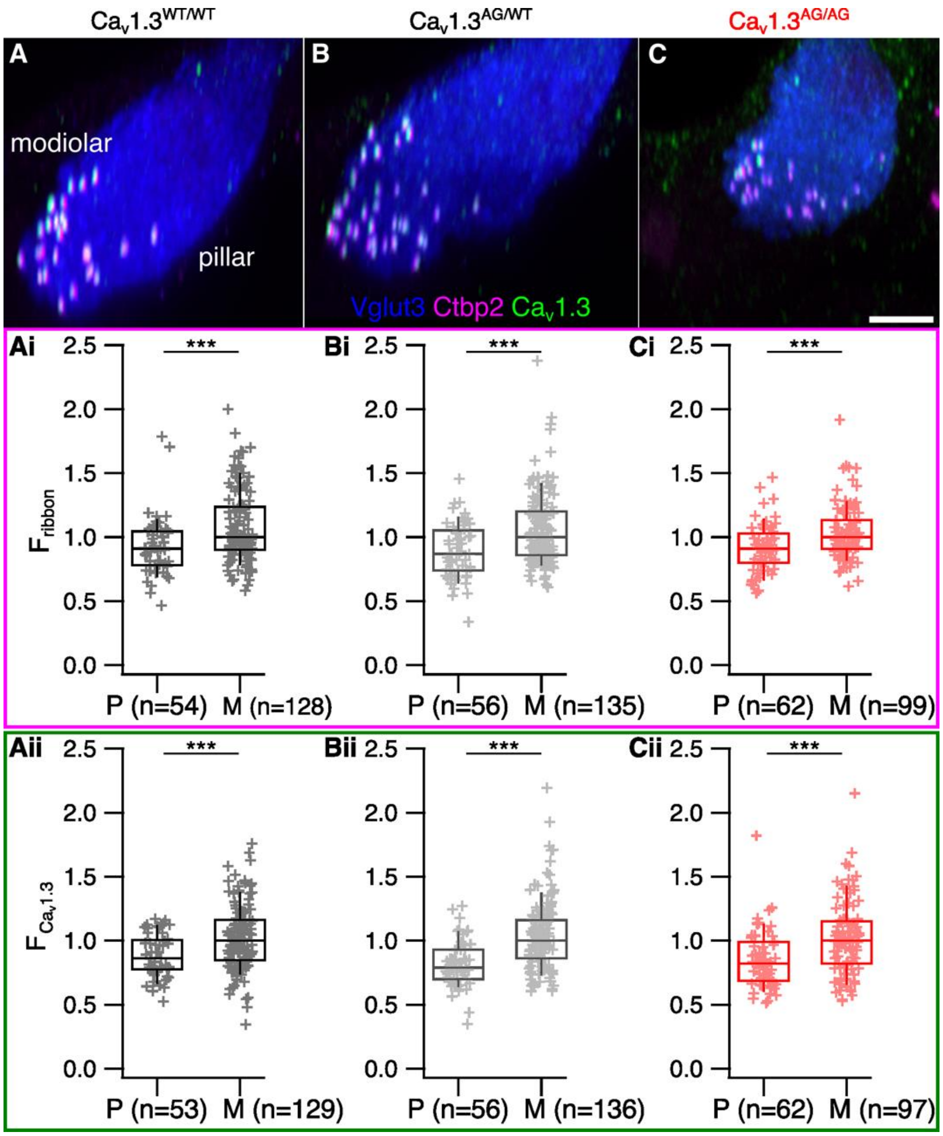


**Fig. S5.**

**Modiolar-pillar gradients of ribbon size and CaV1.3 cluster size are preserved in apical IHCs of CaV1.3$^{AG/WT}$ and CaV1.3$^{AG/AG}$ mice.** (**A-C**) Maximal intensity projections of approximately 2 IHCs from Cav1.3$^{WT/WT}$ (A), Cav1.3$^{AG/WT}$ (B), Cav1.3$^{AG/AG}$ (C) mice immunolabeled against Vglut3, Ctbp2, Cav1.3. (**Ai-Ci**) Comparison of the immunofluorescence intensities of pillar and modiolar ribbons in IHCs of Cav1.3$^{WT/WT}$ (Ai), Cav1.3$^{AG/WT}$ (Bi), Cav1.3$^{AG/AG}$ (Ci) mice. (**Aii-Cii**) Comparison of immunofluorescence intensities of pillar and modiolar Cav1.3 clusters in IHCs of Cav1.3$^{WT/WT}$ (Aii), Cav1.3$^{AG/WT}$ (Bii), Cav1.3$^{AG/AG}$ (Cii) mice. Data were acquired from N = 2 (Cav1.3$^{WT/WT}$), 1 (Cav1.3$^{AG/WT}$), 2 (Cav1.3$^{AG/AG}$) mice. Box-Whisker plots with individual data points overlaid show median, 25$^{th}$ and 75$^{th}$ percentiles (box), 10$^{th}$ and 90$^{th}$ percentiles (whiskers). Statistical significances were determined using two-tailed Wilcoxon rank-sum test for data in (Ai-Ci) and (Aii-Cii). Significances are reported as ***$p < 0.001$.

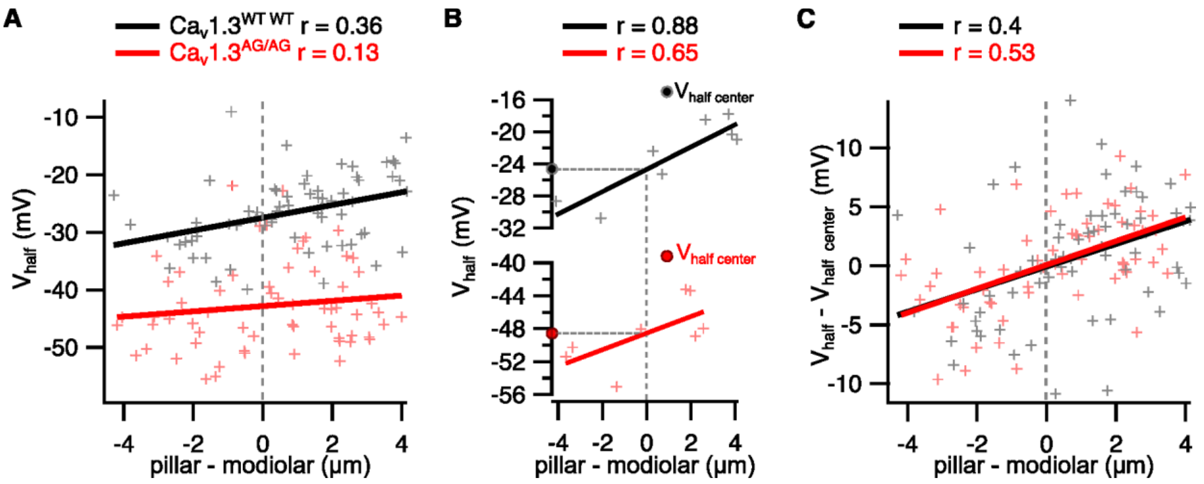


**Fig. S6.**

**Individual IHCs show a pillar-modiolar gradient of voltage of half maximal activation.** (**A**) $V_{half}$ of $Ca^{2+}$ channels at single AZs plotted against their position along the pillar-modiolar axis of the IHC. Thick lines show linear regression lines. Dotted lines indicate the center of the pillar-modiolar axis. (**B**) Same as (A) but from 2 representative cells. $V_{half\ center}$ obtained from the linear fit of $V_{half}$ vs pillar-modiolar position, shows predicted $V_{half}$ of each cell at the center of pillar-modiolar axis. (**C**) $V_{half\ center}$ of each IHC was subtracted from $V_{half}$ of each AZ obtained from the cell, afterwards the data from all the recorded cells were pooled and fitted with linear function. Pearson's correlation coefficient is shown as (r).

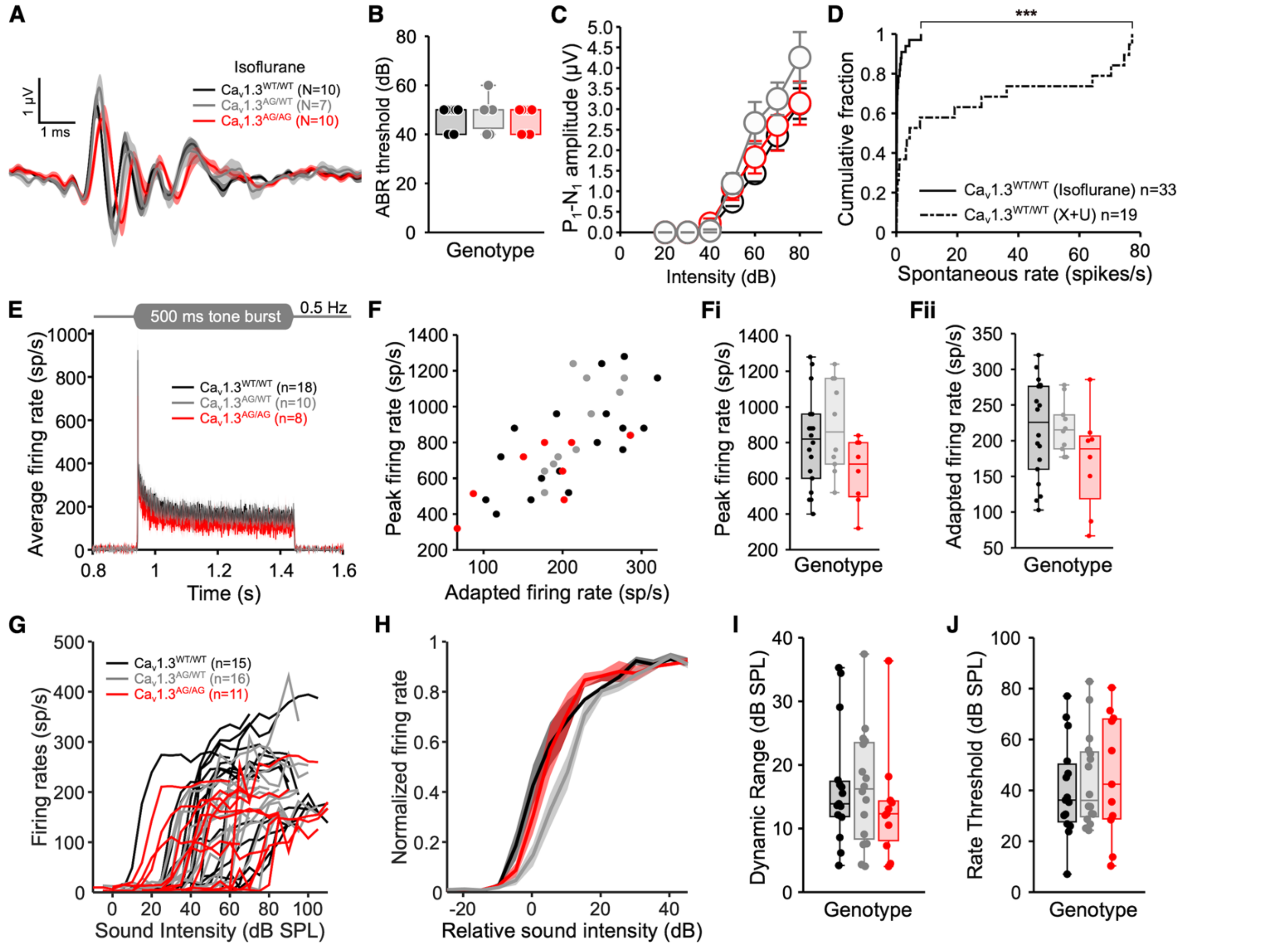


**Fig. S7.**

**Comparable dynamic range of sound encoding by SGNs of Ca$_V$1.3$^{WT/WT}$, Ca$_V$1.3$^{AG/WT}$ and Ca$_V$1.3$^{AG/AG}$ mice.** (**A**) Average ABR waveforms in response to 80 dB clicks recorded in mice under isoflurane anesthesia. (**B**) ABR thresholds in response to click stimuli are comparable in Cav1.3$^{WT/WT}$, Cav1.3$^{AG/WT}$ and Cav1.3$^{AG/AG}$ mice. (**C**) P$_1$-N$_1$ amplitude across different sound levels are comparable in Cav1.3$^{WT/WT}$, Cav1.3$^{AG/WT}$ and Cav1.3$^{AG/AG}$ mice. (**D**) SRs of SGNs recorded from Cav1.3$^{WT/WT}$ mice under isoflurane anesthesia are lower than those recorded under urethane/xylazine aesthesia. (**E**) Average PSTH in response to 500 ms tone burst stimulation at the CF, 30 dB above the threshold level and 0.5 Hz stimulation rate. Shaded areas show ± SEM. (**F, Fi** and **Fii**) Onset (Fi) and adapted (Fii) firing rates calculated from PSTH in (E) are not changed in Cav1.3$^{AG/WT}$ and Cav1.3$^{AG/AG}$ mice. (**G**) Rate level functions (RLFs) of individual SGNs recorded in response to 50 ms stimulation at the CF, 30 dB above the threshold level and stimulation rate of 5 Hz. (**H**) Average and normalized RLFs of SGNs, whereby the RLF of each SGN was further adjusted relative to its threshold (determined from the RLF). Shaded areas show ± SEM. (**I** and **J**) dynamic ranges (I) and the thresholds (J) calculated from RLFs are not changed in Cav1.3$^{AG/WT}$ and Cav1.3$^{AG/AG}$ mice. Single unit recordings were obtained from N = 6 (Cav1.3$^{WT/WT}$), 3 (Cav1.3$^{AG/WT}$), 6 (Cav1.3$^{AG/AG}$) mice. Box-Whisker plots with individual data points overlaid show median, 25$^{th}$ and 75$^{th}$ percentiles (box), and the range (whiskers). Statistical significances were determined using Kruskal-Wallis test for (B), (Fi), (Fii), (I) and (J), Kruskal-Wallis test followed by Tukey-Kramer multiple comparison test for each sound level for (C), Kolmogorov-Smirnov and two-tailed Wilcoxon rank-sum test for (D). Significances are reported as ***$p < 0.001$.

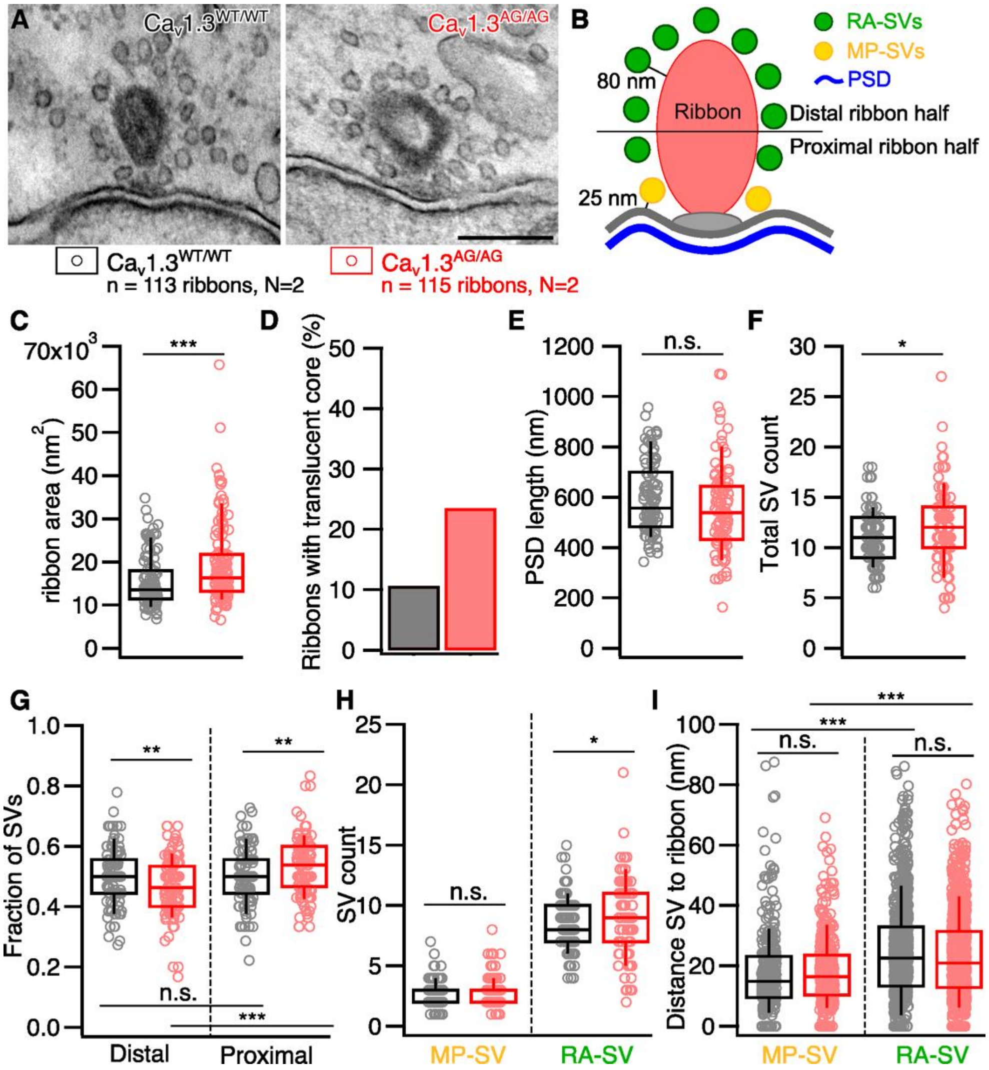


**Fig. S8.**

**Larger fraction of ribbons with hollow cores in Ca$_V$1.3$^{AG/AG}$ IHCs.** (**A**) Representative electron micrographs of the ribbon synapses from Cav1.3$^{WT/WT}$ and Cav1.3$^{AG/AG}$ IHCs. (**B**) Schematic illustration of the quantitative analysis of random EM sections. (**C**) Increased ribbon area in Cav1.3$^{AG/AG}$ IHCs. (**D**) higher percentage of ribbons with translucent core in Cav1.3$^{AG/AG}$ IHCs. (**E**) The length of the postsynaptic density (PSD) is not changed at the afferent synapses of Cav1.3$^{AG/AG}$ IHCs. (**F**) Total vesicle number associated with the ribbons is increased in Cav1.3$^{AG/AG}$ IHCs. (**G**) The fraction of the vesicles associated with the distal half of the ribbon (away from the plasma membrane) is decreased, while those associated with the proximal ribbon half (close to the plasma membrane) is increased in Cav1.3$^{AG/AG}$ IHCs. (**H**) The number of the membrane proximal synaptic vesicles is unchanged, while the number of the ribbon associated vesicles is increased in Cav1.3$^{AG/AG}$ IHCs. (**I**) The distance of the membrane proximal and ribbon associated vesicles from the ribbon is not changed in Cav1.3$^{AG/AG}$ IHCs. Each genotype represents data from N = 2 mice. Box-Whisker plots with individual data points overlaid show median, 25$^{th}$ and 75$^{th}$ percentiles (box), 10$^{th}$ and 90$^{th}$ percentiles (whiskers). Statistical significances were determined using two-tailed Wilcoxon rank-sum test for (C), (E), (F), (H) and (I) and two-tailed t-test for (G)**.** Significances are reported as *$p < 0.05$, **$p < 0.01$, ***$p < 0.001$.

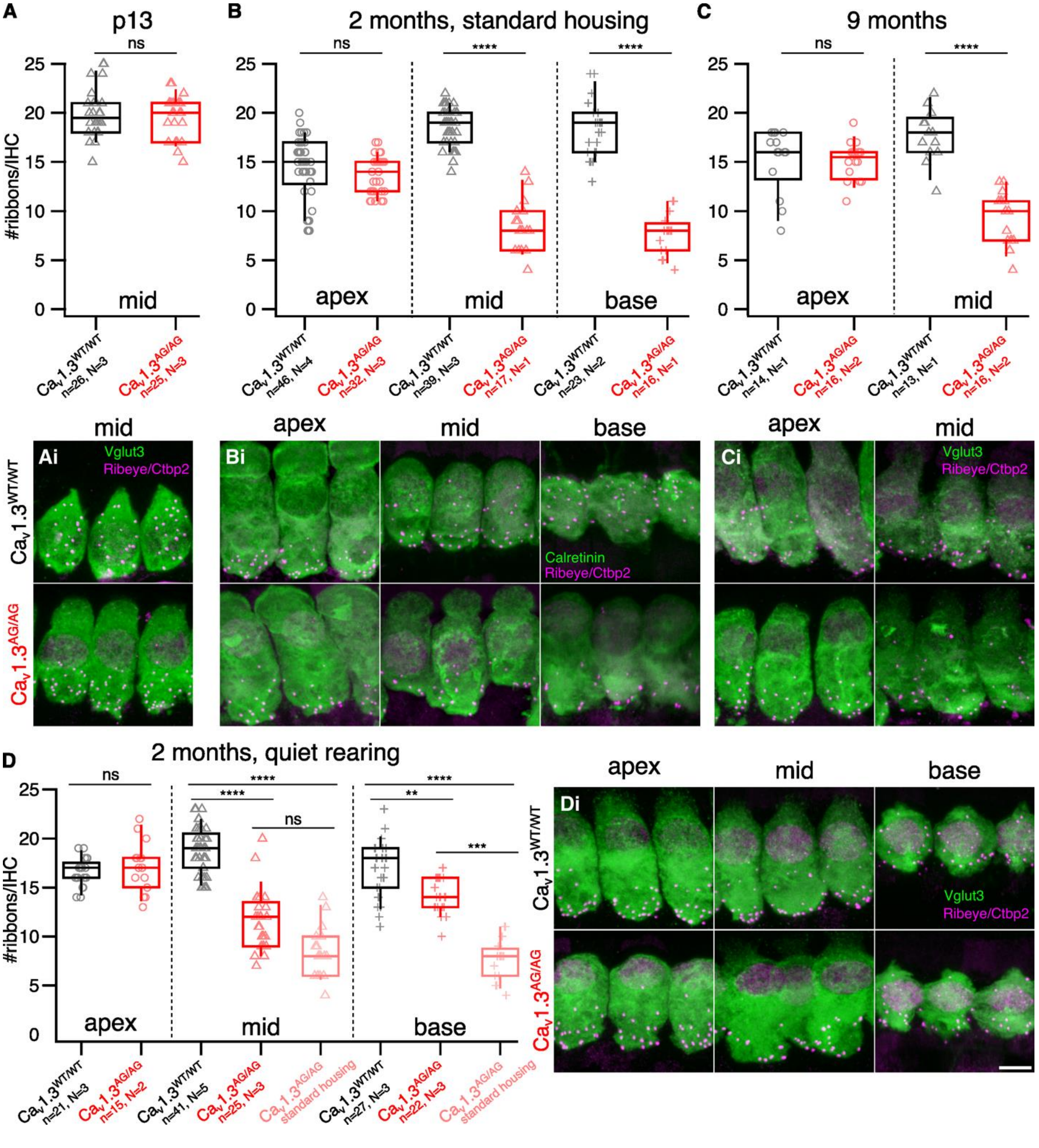


**Fig. S9.**

**Ca$_V$1.3$^{AG/AG}$ IHCs exhibit loss of ribbons at the middle and basal turns of the cochlea after the hearing onset.** (**A-D**) Number of the ribbons at different tonotopic locations was counted in the confocal stacks of IHCs of p13 (A), 2-month-old (B), 9-month-old (C) and 2-month-old quietly reared (D) mice. (**Ai-Di**) Maximal intensity projections of the representative IHCs immunolabeled against Ribeye/Ctbp2 and calretinin or Vglut3. Scale bar = 5 µm. Box-Whisker plots with individual data points overlaid show median, 25$^{th}$ and 50$^{th}$ quartiles (box), 10$^{th}$ and 90$^{th}$ percentiles (whiskers). Statistical significances were determined using two-tailed Wilcoxon rank-sum test for each tonotopic location followed by Bonferroni-Holm multiple comparison correction for (A), (B) and (C) and Kruskal-Wallis followed by Dunn's multiple comparison test for middle and basal cochlear regions for (D). Significances are reported as ***$p < 0.001$, ****$p < 0.0001$.

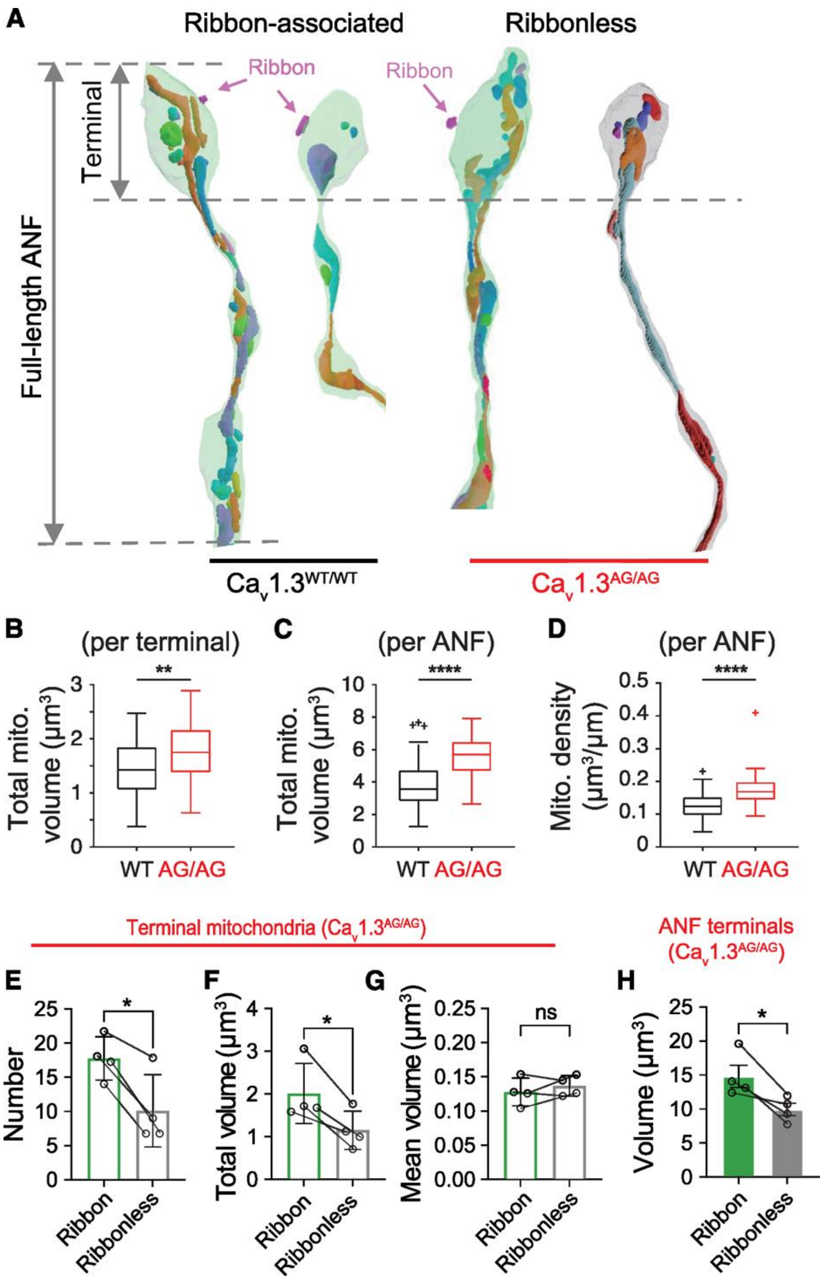


**Fig. S10.**

**Quantification of SGN mitochondria content in SBEM reconstructions of $Ca_V1.3^{WT/WT}$ and $Ca_V1.3^{AG/AG}$ mid-cochlear region.** (**A**) Mitochondrial reconstructions of example auditory nerve fiber (ANF, peripheral neurite of SGN) postsynaptic to the $Ca_v1.3^{WT/WT}$ (left) and $Ca_v1.3^{AG/AG}$ (right) IHCs. (**B** and **C**) In contrast to $Ca_v1.3^{WT/WT}$ (black), total mitochondrial volumes are larger in both terminals (b) and full-length peripheral neurites (c) of ribbon-associated SGNs in $Ca_v1.3^{AG/AG}$ (red) cochlea. (**D**) Higher ANF mitochondrial density of $Ca_v1.3^{AG/AG}$ (red) than that of $Ca_v1.3^{WT/WT}$ cochlea (black). (**E** and **F**) For SGNs on $Ca_v1.3^{AG/AG}$ IHCs, the ribbon-associated terminal (green) features a greater number and larger total volume of mitochondria than those without a ribbon (grey). (**G**) Mean sizes of mitochondria are comparable between ribbon-associated and ribbonless terminals. (**H**) Ribbon-associated SGNs (green) have a larger terminal size than ribbonless SGNs (grey). Each genotype represents data from N = 2 mice. Statistical significances were determined using two-tailed t-test for (B), (C) and (D) and paired t-test for (E), (F), (G) and (H). Significances are reported as *$p < 0.05$, **$p < 0.01$, and ****$p < 0.0001$.

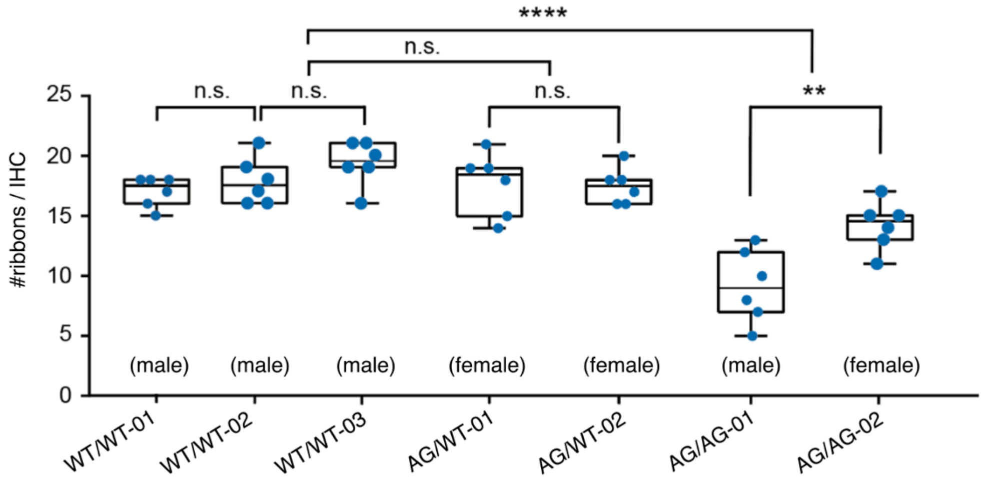


**Fig. S11.**

**Quantification of IHC synaptic ribbon numbers in SBEM reconstructions of mid-cochlear regions in individual $Ca_V1.3^{WT/WT}$, $Ca_V1.3^{AG/WT}$ and $Ca_V1.3^{AG/AG}$ animals**. The number of the ribbons is maintained in $Ca_V1.3^{AG/WT}$ IHCs and is reduced in $Ca_V1.3^{AG/AG}$ IHCs. Box-Whisker plots with individual data points overlaid show median, 25$^{th}$ and 75$^{th}$ percentiles (box), and the range (whiskers). Statistical significances were determined using two-tailed t-test. Significances are reported as $^{**}p < 0.01$, $^{****}p < 0.0001$.

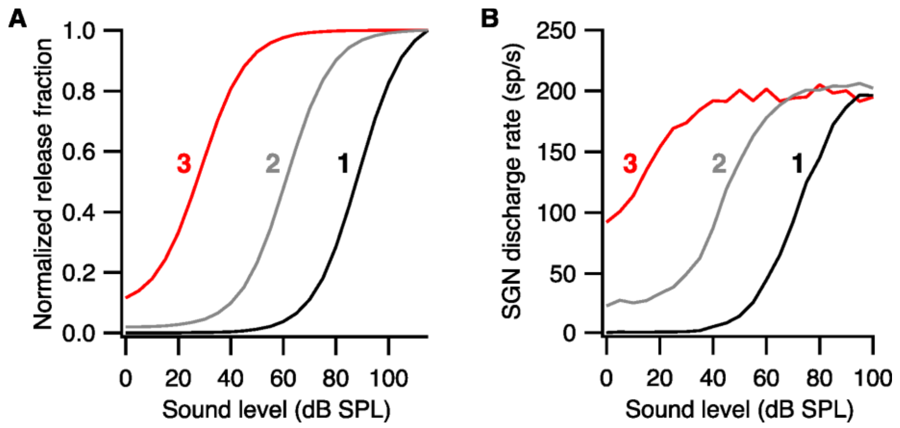


**Fig. S12.**

**The effects of modifying model parameters A and B on the SGN rate-level function.** (**A** and **B**) Normalized release fractions (A) and their corresponding rate-level functions (B). The exact parameters can be found in Table S1.

**Table S1.**

**Model parameters.** Parameters A and B of the release fraction equation were modified, but the rest of the parameters were kept constant.

| **Parameters** | **1** | **2** | **3** |
|---|---|---|---|
| A | 1 | 1 | 0.1 |
| B | 1200 | 50 | 1.1 |
| g | 1660 | 1660 | 1660 |
| dt (s) | 0.0001 | 0.0001 | 0.0001 |
| y (replenishment rate) | 16.6 | 16.6 | 16.6 |
| l (rate of loss from the cleft) | 500 | 500 | 500 |
| r (rate of return from the cleft) | 12500 | 12500 | 12500 |
| x (rate of release from reprocessing to free transmitter) | 3000 | 3000 | 3000 |
| h (firing probability scaling factor) | 10000 | 10000 | 10000 |
